# Cytotoxic CD4+ exTregs expressing KLRG1 retain immunosuppressive function in cancer

**DOI:** 10.64898/2026.11.09.765461

**Authors:** Collin Jugler, Kade R. Copple, Jessica C. Hill, Phuong Nguyen, Mouneeb M. Choudry, Jarl F. Carnahan, Nilika Bhattacharya, Fan Shen, Niharika Abbaraju, Jessica N. Lancaster, Feda H. Hamdan, Yousef Zakharia, Jack R. Andrews, Paul E. Andrews, Haidar M. Abdul-Muhsin, Casey R. Ager

## Abstract

The role of cytotoxic CD4+ T cells in cancer remains controversial, with both pro- and anti-tumor functionality described. Based on a prior observation that putatively cytotoxic KLRG1+ FoxP3-conventional T cells (Tconv) associated with tumor progression, we deeply investigated the phenotype and function of KLRG1+ Tconv in multiple murine tumor models and in human clear cell renal cell carcinoma (ccRCC) specimens. We found KLRG1+ Tconv have a conserved, highly activated, terminally differentiated phenotype across species. In murine tumors, KLRG1+ Tconv resemble tumor-infiltrating regulatory T cells (Tregs) at both transcriptional and proteomic levels, upregulating a spectrum of exhaustion-related checkpoint receptors, producing IL-10 and granzyme B (GzmB), and silencing expression of IFNγ and TNFα. Functionally, KLRG1+ Tconv exhibited immunosuppressive function in *ex vivo* assays. Using an *in vivo* Treg fate mapping approach in tumor-bearing mice, we found a subset of KLRG1+ Tconv are derived from Tregs that have lost FoxP3 protein expression. *Ex vivo* functional assays demonstrated the exTreg component of the KLRG1+ Tconv compartment is responsible for IL-10 production and reduced pro-inflammatory cytokine expression. In treatment studies, we found that KLRG1-targeted antibody-mediated depletion was more efficient than anti-KLRG1 blockade at slowing tumor growth and improving survival, with responses associating with depletion of intratumoral KLRG1+ Tconv and Tregs. In sum, our study discovered a novel conserved pathway of CD4+ T cell plasticity, wherein intratumoral, cytotoxic KLRG1+ CD4+ Tconv cells can be derived from Tregs and maintain immunosuppressive function, despite loss of FoxP3, and that KLRG1-targeted depletion may outperform KLRG1 blockade as an immunotherapeutic strategy in cancer.

**One sentence summary:** KLRG1-expressing FoxP3-negative CD4+ T cells in tumors harbor a pool of exTregs that retain an immunosuppressive and cytotoxic profile.

## INTRODUCTION

CD4+ T cells play a central role in shaping anti-tumor immunity, however the individual contributions of various CD4+ T cell subsets to this process remain unclear. Conventional CD4+ T cell (Tconv) effector and memory subsets possessing “helper” function, counterbalanced by immunosuppressive and tumor-promoting FoxP3+ regulatory T cells (Tregs) are well-described (*1, 2*). In contrast, the etiology and roles of non-canonical CD4+ populations in the tumor microenvironment (TME) including FoxP3-tolerogenic CD4+ (Tr1 cells) and cytotoxic (CD4+ CTLs) CD4+ T cells remain controversial (*3–8*). An additional complicating factor in the study of these subsets is the well-recognized plasticity or lineage instability of CD4+ T cells, particularly in complex tissue environments such as the TME (*5, 9, 10*). For example, in line with the key tenet of peripheral tolerance, Tconv can be converted into FoxP3+ Tregs and contribute to direct antigen-specific suppression of antitumor immunity (*11, 12*). Conversely, Tregs can lose FoxP3 expression under certain conditions and gain inflammatory antitumor functionality (*13–15*). How this plasticity shapes the etiology or function of non-canonical CD4+ subsets (e.g., Tr1, CTL, etc.) remains obscure and warrants further investigation.

In prior work, we explored the immunological dynamics of cancer immunoediting in a murine prostate cancer model and identified a CD4+ Tconv subset marked by expression of KLRG1 (killer cell lectin-like receptor G1) which was highly associated with immune evasive tumors, was absent in tumors undergoing curative response to immunotherapy, and could be identified with increasing frequency in advanced renal cell carcinomas (*16*). While KLRG1 has well-defined roles in natural killer (NK) cell and CD8+ T cell biology (*17, 18*), the functional role of KLRG1-expressing CD4+ T cells has received limited attention. KLRG1 is an emerging feature of CD4+ CTLs in circulation in patients from paired blood and tumor tissues in bladder cancer and melanoma (*19, 20*). It has also been established that KLRG1+ Tregs are more potent immune suppressors than their KLRG1-counterparts (*21, 22*). Perhaps reflecting this duality, KLRG1-targeted immune checkpoint blockade for the treatment of various cancers has been evaluated and has shown to have modest monotherapy efficacy which can be improved only in combination with anti-PD-1/L1 blockade in an NK- and/or CD8-dependent manner (*23–25*). Greater understanding of the unique contributions of KLRG1+ CD4+ T cells to the antitumor response may have utility in improving KLRG1-targeted therapeutics and development of novel therapeutic strategies to rationally modulate CD4+ T cell functionality in cancer.

To address these knowledge gaps, we pursued a systematic characterization and functional evaluation of KLRG1+ CD4+ Tconv. We employed high-parameter spectral flow cytometry exceeding 40-50+ parameters to quantify the abundance and phenotypic signatures of KLRG1+ Tconv across several murine tumors spanning distinct genetic strains and levels of immune infiltration, as well as in healthy donor peripheral blood mononuclear cells (PBMCs) and in patients with clear cell renal cell carcinoma (ccRCC). We investigated the cytokine production capacity of this subset and performed *ex vivo* functional assays to determine how tumor-derived KLRG1+ CD4+ Tconv modulate T cell and dendritic cell (DC) function. Using transcriptomic analyses, we map a core transcriptional signature of the intratumoral KLRG1+ Tconv cell state and use *in vivo* fate-mapping approaches to explore potential lineage connections between KLRG1+ CD4+ Tconv and Tregs. To investigate therapeutic implications, we explored the relative efficacy and immune correlates of response to antibodies engineered for KLRG1-targeted depletion versus blockade. In total, our findings reveal a previously unappreciated etiology and immunomodulatory role for KLRG1+ CD4+ Tconv in cancer.

## RESULTS

### The KLRG1+ Tconv phenotype is conserved across murine tumors

Our previous work identified CD4+ FoxP3-KLRG1+ T cells (KLRG1+ Tconv) as a putatively tumor-promoting T cell subset in the NPK-C1 prostate cancer model (*16, 26*). To first determine whether the KLRG1+ Tconv phenotype is a conserved feature of the murine TME, we screened several syngeneic tumor models spanning multiple strains, genders, and levels of sensitivity to immunotherapy for the presence of this subset. Specifically, we established C57BL/6-syngeneic B16-F10 (melanoma), MB49 (bladder), MCA205 (fibrosarcoma), or MC38 (colon) tumors, BALB/c-syngeneic RENCA (renal) tumors, or FVB-syngeneic MycCaP (prostate) tumors in male and/or female mice as appropriate (see Methods for model-specific gender distribution), then performed 42-color spectral flow cytometry on freshly dissociated tumors when tumors reached an approximate average volume of at least 150 mm^3^ (Figure 1A). We employed an established semi-supervised FlowSOM clustering and PaCMAP embedding analysis pipeline (*16, 26, 27*) to analyze these data, first delineating major tumor-infiltrating immune lineages based on canonical lineage-defining immune marker expression (Figure 1B-C). Tumor immune composition was heterogeneous across models, as expected (Figure 1D). Focusing exclusively on the CD4+ FoxP3-Tconv compartment, our sub-clustering revealed 11 distinct clusters with clear meta-clusters defining naïve-like, KLRG1+, tissue-resident memory (TRM), and heterogeneous effector-like states (Figure 1E-F). The uniquely clustered KLRG1+ population was detected in all models, with highest relative proportionality (approximately 10-30% among CD4+ Tconv) in MB49, MCA205, and MC38 models, and a lower relative proportionality (approximately 1-10%) in immunologically “colder” models including B16-F10, MycCaP, and RENCA models (Figure 1G). As a proportion of total tumor-infiltrating immune cells, MCA205 exhibits the largest frequency of KLRG1+ Tconv cells (*p*<0.0001 across all other models). These data support the notion that KLRG1+ Tconv cells are a conserved population of the mouse TME across strain and gender and may be relatively enriched in highly immune-infiltrated tumors.

**Figure 1:**
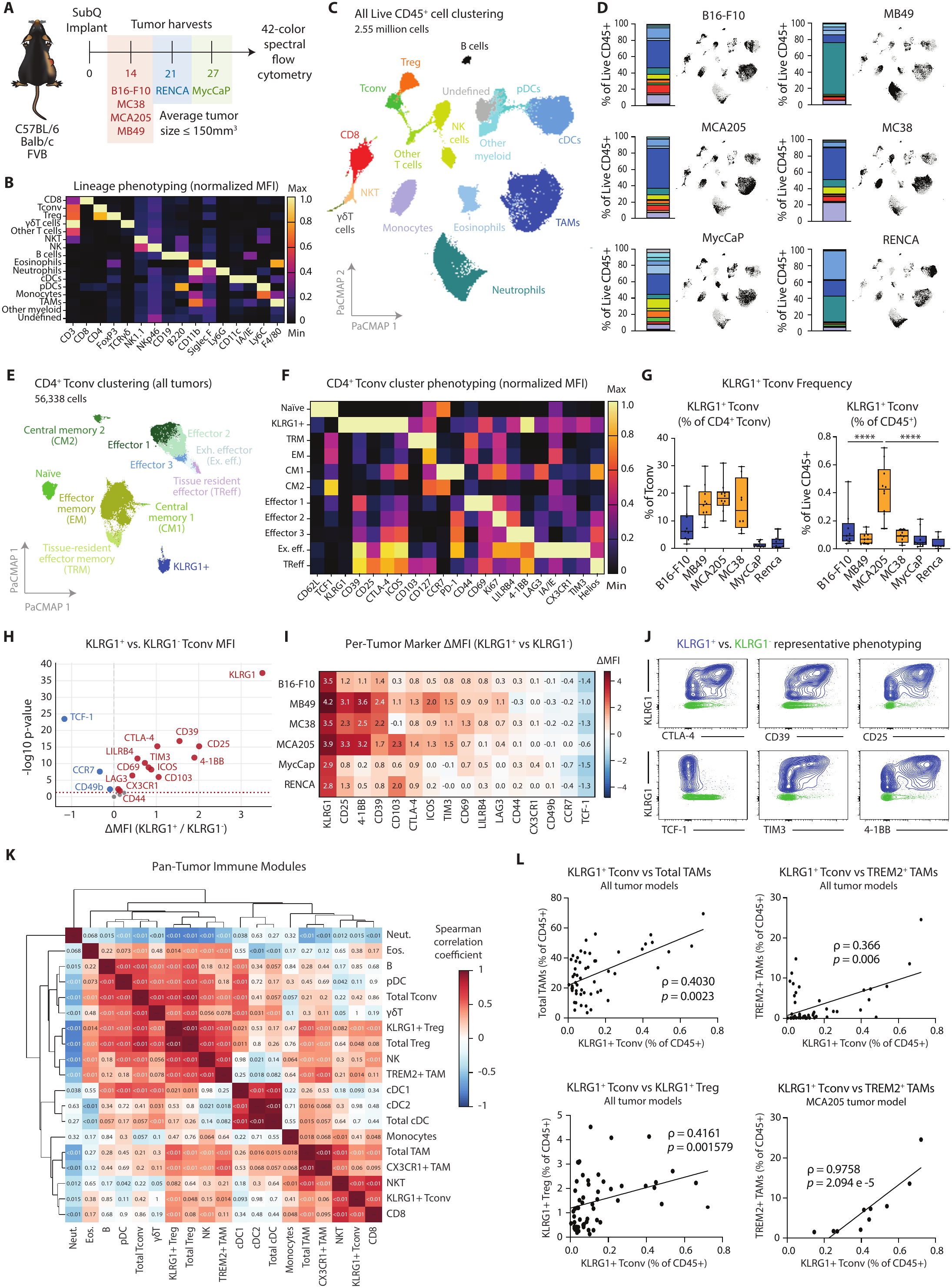
A terminally differentiated KLRG1+ Tconv phenotype is conserved across murine tumor models. (**A**) Conceptual schematic of immune profiling timeline. (**B**) Normalized heatmap of lineage-defining marker MFI of all live CD45+ cells across all tumor models. (**C**) PaCMAP plot of live CD45+ cells across all tumor models. (**D**) Each model overlaid on the PaCMAP plot with the associated immune composition of clusters in C. (**E**) PaCMAP plot of the CD4+ Tconv cluster from C. (**F**) Normalized heatmap of phenotyping marker MFI in CD4+ Tconv across all tumor models. (**G**) Quantitation of KLRG1+ Tconv as a fraction of CD4+ cells (left) or as a fraction of all live immune cells (right). Warm colors denote “hot” tumor models, cool colors denote “cold” tumors. Ordinary one-way ANOVA with Holm-Sidak correction was performed. (**H**) Delta arcsinh-transformed (cofactor = 1500) for paired MFI values for each marker in KLRG1+ or KLRG1-Tconv across all tumor models. Two-sided paired t-test with Benjamini-Hochberg (BH) correction was performed. Red is up in KLRG1+ Tconv, blue is up in KLRG1-Tconv. (**I**) Delta MFIs for each significant marker from H within each tumor model. Cell color indicates whether the delta MFI is positive (red) or negative (blue). (**J**) Manual phenotyping of select significant markers from H and I. (**K**) Correlation matrix of frequencies of immune populations frequencies from all tumor models. Spearman correlation was performed for each relationship and *p*-values adjusted for BH correction. Cell color indicates Spearman rho value and cell values are the adjusted *p*-value for the correlation. (**L**) Targeted correlations of KLRG1+ Tconv against populations of interest. Spearman rho is listed above the uncorrected p-value for that correlation plot. All clustering is derived from FlowSOM. Data shown are *n* = 8-10 with approximately equal numbers male and female mice where the tumor model permits and are a representative replicate out of two independent immune profiling experiments across multiple tumor models. **** = *p* ≤ 0.0001

### KLRG1+ Tconv are terminally differentiated and correlate with tumor macrophage infiltration

To define the phenotype of KLRG1+ Tconv in tumors, we leveraged our 42-parameter spectral flow cytometry panel to extract the pan-cancer phenotypic profile of KLRG1+ Tconv cells in our 6-model dataset. Quantifying the relative mean fluorescent intensity (MFI) of each phenotypic marker between the KLRG1+ and KLRG1-Tconv fractions across all models, we observed that the KLRG1+ phenotype in mice is defined by significant upregulation of CD25, 4-1BB, CD39, CD103, CTLA-4, ICOS, TIM3, CD69, LILRB4, LAG3, CX3CR1, and CD44 with reduced expression of TCF-1, CCR7, and CD49b (Figure 1H-J). This signature was largely conserved when individually profiling KLRG1+ Tconv in each individual model, with minor discrepancies in relative expression of CD103 noted in B16-F10 and MC38 (Figure 1I). These findings indicate KLRG1+ Tconv in tumors exhibit a conserved phenotype marked by an activated and/or terminally differentiated state, in line with existing literature (*17, 21, 22, 28*). To further explore potential functional roles of KLRG1+ Tconv in the TME, we next asked whether KLRG1+ Tconv is associated with recurrent cellular programs across tumors. For this, we employed a Spearman correlation analysis comparing the relative frequencies of all defined immune lineages across tumors followed by hierarchical clustering. The resultant correlation matrix (Figure 1K-L) revealed statistically significant positive correlations between KLRG1+ Tconv and CD8+ T cells (r=0.565; adj. *p*<0.01), NKT cells (r=0.655; adj. *p*<0.01), total tumor-associated macrophages (TAMs) (r=0.403; adj. *p*<0.01), TREM2+ TAMs (r=0.366; adj. *p*=0.014), Tregs (r=0.305; adj. *p*=0.048), and KLRG1+ Tregs (r=0.416; adj. *p*<0.01). The correlation between KLRG1+ Tconv and TREM2+ TAMs was particularly enriched in MCA205 tumors that possess a relatively high abundance of KLRG1+ Tconv (Figure 1L). These descriptive findings support the hypothesis that KLRG1+ Tconv associate with an immunosuppressive program spanning myeloid, cytotoxic, and CD4+ T cell compartments across tumors.

### KLRG1+ Tconv in humans maintain a cytotoxic TEMRA phenotype

Having interrogated the phenotype of KLRG1+ Tconv in murine models, we next sought to test whether KLRG1+ Tconv could be identified in human tumors and investigate to what extent the phenotypic determinants of this population are conserved in humans. To accomplish this, we leveraged an in-house optimized 51-color spectral flow cytometry panel applied on fresh surgical samples from 5 patients with ccRCC who received standard of care nephrectomy resections at Mayo Clinic Arizona (Figure 2A; S1A). High dimensional flow cytometric analysis of the tumor tissue alongside cryopreserved healthy volunteer PBMCs resulted in clearly defined and distinct immune lineages, with expected differences in the immune composition between tumor and PBMC compartments, marked by neutrophil and CD8+ T cell populations expanding proportionally in ccRCC tissue, whereas B cells and monocytes predominate in PBMCs (Figure 2B-C; S1B). We performed an additional round of sub-clustering on the FoxP3-CD4+ Tconv cluster, which yielded 13 subclusters. We found expression of KLRG1 within three clusters, with greatest relative MFI in the KLRG1+ 1 cluster where KLRG1 appeared to be a defining marker alongside GzmB and CX3CR1, as observed in our murine data. This cluster was present in both PBMCs and tumors, marking approximately 33.42% and 12.67% of the CD4+ Tconv compartment, respectively (Figure 2D-F). We next tested whether KLRG1+ Tconv were enriched for a terminally differentiated, cytotoxic phenotype. We observed a trending enrichment for GzmB+CD57+ enrichment in KLRG1+ Tconv relative to the KLRG1-Tconv counterpart in both PBMCs and ccRCC tissue (Figure 2G-H). To define a phenotypic signature of KLRG1+ Tconv using unsupervised approaches, we compared the relative MFI of all markers in our panel between KLRG1+ and KLRG1-Tconv, performing independent analyses for PBMCs and ccRCC tumors. In PBMCs, KLRG1+ Tconv expressed higher levels of 4-1BB, CXCR3, and CXCR5 than KLRG1-Tconv. In tumors, KLRG1+ Tconv trended towards higher expression of CD127, CD45RA, CX3CR1, and PD-L2, with reduced expression of CD39, CCR7, CCR6, CD304 and CD27 relative to KLRG1-Tconv (Figure 2I; S1C), however only CD39 remained significant after multiple testing correction. The differential levels of CD45RA and CCR7 led us to hypothesize that the terminally differentiated KLRG1+ Tconv may also have a TEMRA phenotype. Indeed, we found that KLRG1+ Tconv in ccRCC tumors were trending for an enrichment in a CCR7-CD45RA+ TEMRA phenotype versus KLRG1-Tconv (p=0.0625; Figure 2J). In total, this data supports a conserved phenotype of KLRG1+ Tconv between mouse and human tumors as human KLRG1+ Tconv tend to resemble terminal effectors enriched in a TEMRA phenotype in RCC tumors and are marked by high levels of granzyme B and CX3CR1, suggestive of late differentiation and putative cytotoxic potential.

**Figure 2:**
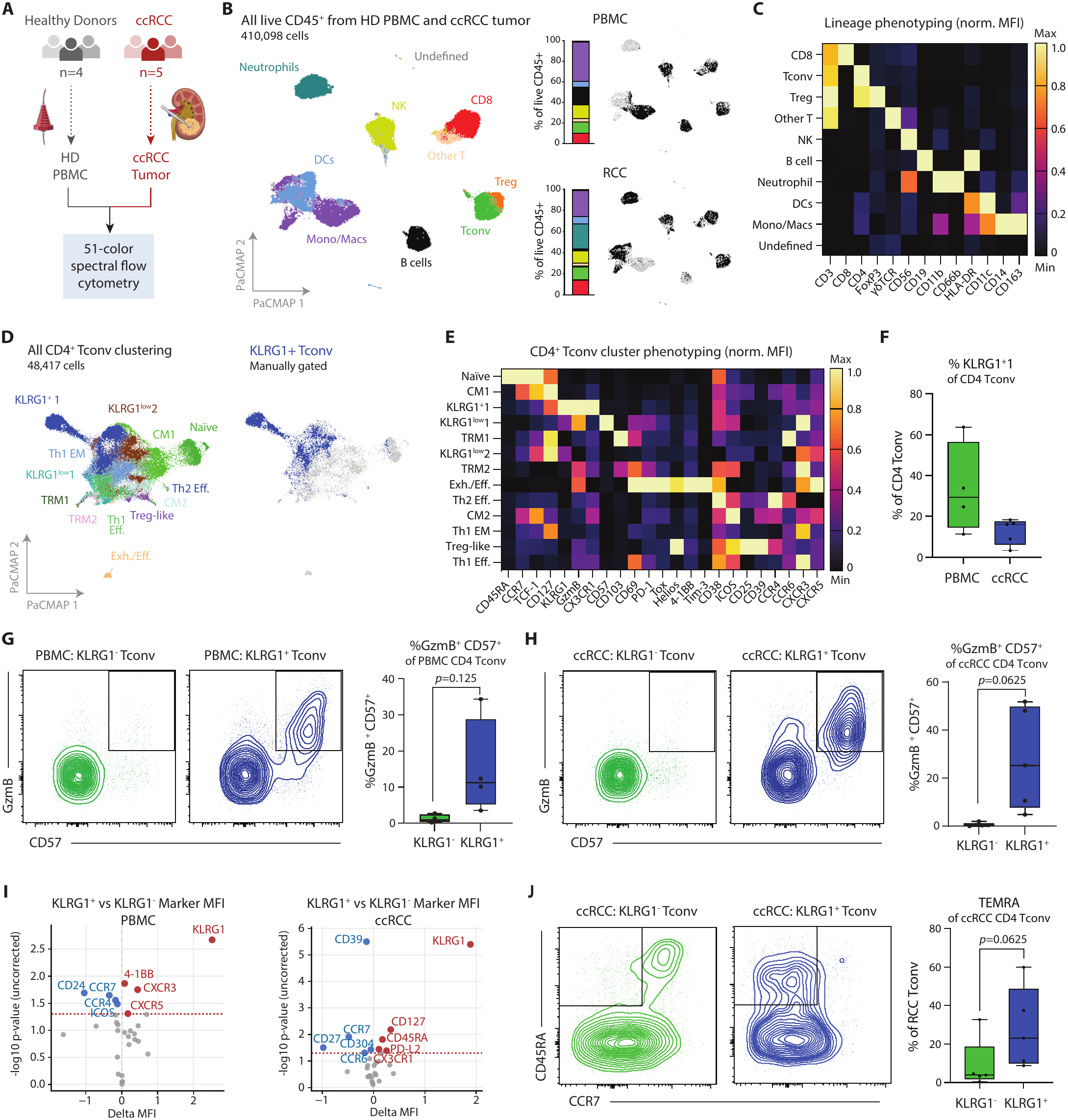
KLRG1+ Tconv in humans maintain a terminally differentiated, cytotoxic, TEMRA phenotype. (**A**) Conceptual schematic of tissue acquisition and experimental design. HD PBMC= healthy donor peripheral blood mononuclear cells; ccRCC = clear cell renal carcinoma. (**B**) PaCMAP of live CD45+ cells from HD PBMCs and ccRCC tissue. Clusters are derived from FlowSOM (left); HD PBMC samples (right top) or ccRCC samples (right bottom) overlaid on the PaCMAP with the quantified immune composition. (**C**) Normalized heatmap of lineage-defining marker MFI across all live CD45+ cells from both HD PBMC and ccRCC samples. (**D**) PaCMAP plot of the CD4+ Tconv cluster from B. Clusters are derived from FlowSOM (left) and manually gated KLRG1+ Tconv are overlaid on the PaCMAP (right). (**E**) Normalized heatmap of phenotyping marker MFI across all CD4+ Tconv from both HD PBMC and ccRCC samples. (**F**) Quantitation of the frequency of the KLRG1+ 1 cluster from D and E in HD PBMC and ccRCC tissue. (**G**) Representative plots and quantitation of GzmB+ CD57+ Tconv from HD PBMCs. Wilcoxon matched pairs signed rank test was used for statistical testing. PBMCs *n* = 4. (**H**) Representative plots and quantitation of GzmB+ CD57+ Tconv from ccRCC tissue. Wilcoxon matched pairs signed rank test was used for statistical testing. ccRCC *n* = 5. (**I**) The delta arcsinh-transformed (cofactor = 1500) paired MFI values for each marker from KLRG1+ versus KLRG1-Tconv from HD PBMC (left plot) or from ccRCC tissue (right plot). MFI values were averaged across samples from the respective group. Two-sided paired t-test was performed without correcting *p*-values for multiple testing. (**J**) Representative plots and quantitation of TEMRA Tconv from ccRCC tissue. Wilcoxon matched pairs signed rank test was used for statistical testing. ccRCC *n* = 5

### Tumor-infiltrating KLRG1+ Tconv are functionally immunosuppressive

To define the functional characteristics of tumor-infiltrating KLRG1+ Tconv, we first investigated their cytokine profile by *ex vivo* restimulation and intracellular cytokine staining (ICS) in murine models (Figure 3A; S2A). For this, we employed two tumor models: (i) the NPK-C1 model in which we first defined the KLRG1+ Tconv phenotype(*16*), and (ii) the MCA205 model, which we found possessed the largest KLRG1+ Tconv fraction of all models we profiled (Figure 1G). We harvested established tumors (>200mm^3^) of each type, dissociated, then performed polyclonal *ex vivo* restimulation via PMA/ionomycin and subsequent ICS. We found KLRG1+ Tconv from both NPK-C1 and MCA205 exhibited enhanced expression of GzmB (*p*=0.031, *p*=0.0078) and IL-10 (*p*=0.031, *p*=0.0078), and reduced expression of IFNγ (*p*=0.031, *p*=0.0078) and TNFα (*p*=0.031, *p*=0.0078) as compared to their KLRG1-Tconv counterparts (Figure 3B-C). In this assay, the KLRG1+ Tconv compartment resembled the cytokine profile of KLRG1+/- Tregs, suggesting tumor-infiltrating KLRG1+ Tconv may be functionally immunosuppressive. To test whether KLRG1+ Tconv are functionally suppressive, we flow sorted KLRG1+ vs KLRG1-Tconv and Tregs from MCA205 tumors using *Foxp3*^RFP^ reporter mice for *in vitro* T cell micro-suppression assays (Figure 3D). We observed that co-culture with KLRG1+ Tconv led to a significant drop in responder T cell division as compared to co-culture with KLRG1-Tconv (*p*=0.0023); this magnitude of T cell suppression was equivalent to that seen by tumor-derived KLRG1+ or KLRG1-Tregs (Figure 3F). These data support the notion that KLRG1+ Tconv are sufficient to suppress T cell proliferation. Related findings described a cytotoxic CD4+ T cell expressing IL-10 generated via neoantigen peptide vaccination that suppressed antitumor immunity by cytotoxic removal of conventional type 1 dendritic cells (cDC1s) (*6*). To test whether KLRG1+ Tconv manipulate cDC1 viability or phenotype, we sorted MCA205-infiltrating KLRG1+ and KLRG1-Tconv and Tregs for *ex vivo* co-culture with bone marrow derived dendritic cells (BMDCs) followed by spectral flow cytometry to assess BMDC viability and/or licensing (Figure 3G). We did not observe a significant change in DC viability upon co-culture with KLRG1+ Tconv in the presence or absence of tumor antigen (Figure S2B) suggesting MCA205-infiltrating KLRG1+ Tconv may lack cytotoxic function against cDCs. However, while KLRG1-Tconv were effective at licensing cDC activation as measured by upregulation of CD80 (*p*=0.0023 in total cDCs; *p*=0.0012 in cDC1s) and CD86 (*p*=0.0029 in total cDCs; *p*=0.0041 in cDC1s), KLRG1+ Tconv did not license DC activation, phenocopying co-culture with KLRG1+/-Tregs (Figure 3H). Taken together, our findings suggest tumor-infiltrating KLRG1+ Tconv are sufficient to produce both cytotoxic and immunosuppressive cytokines, are functionally immunosuppressive against T cell proliferation, and lack DC licensing capacity.

**Figure 3:**
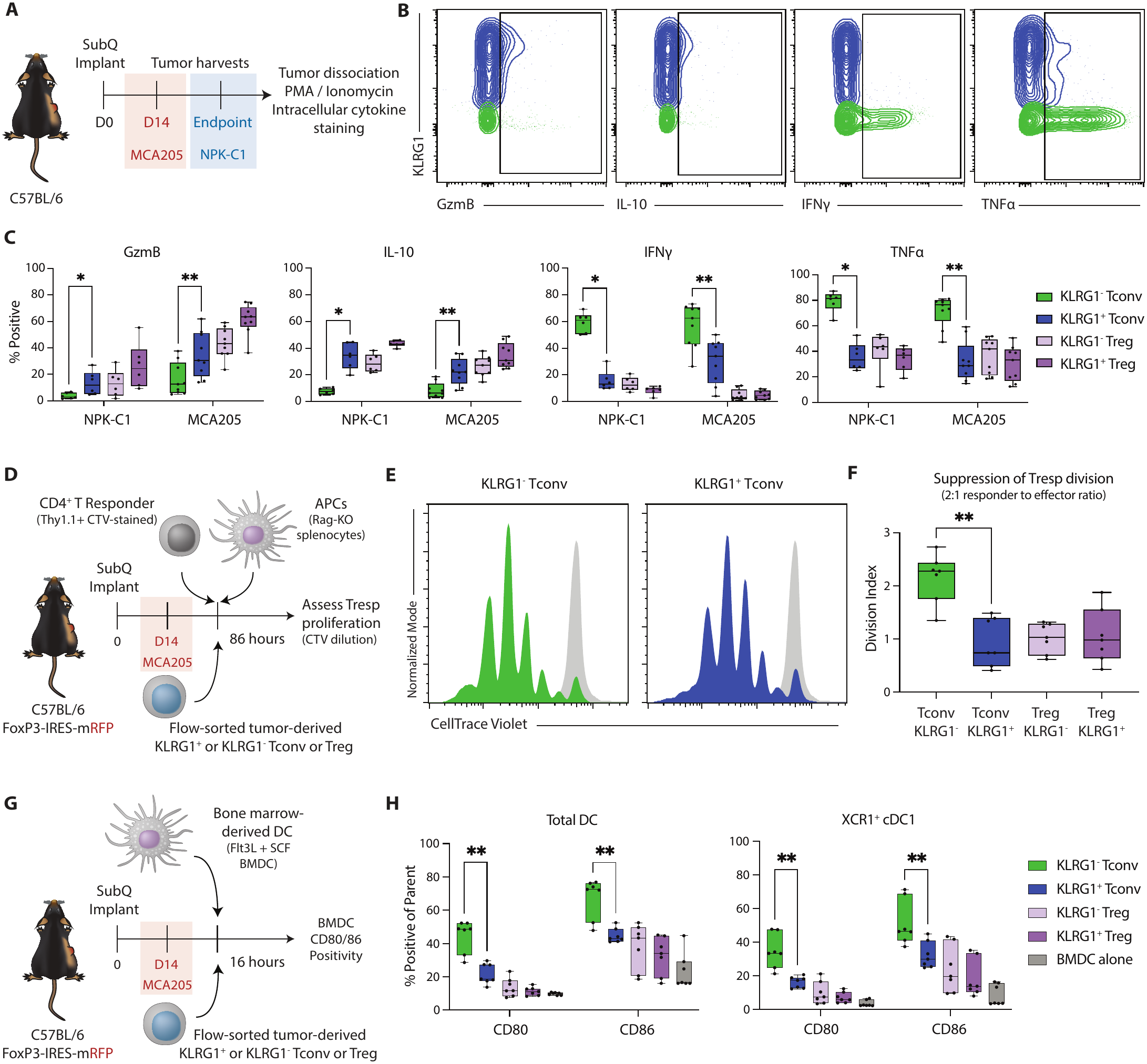
KLRG1+ Tconv cells have suppressive functional characteristics. (**A**) Conceptual schematic of timing for cytokine analysis. (**B**) Representative examples of cytokine gating. Blue contours are gated as KLRG1+ Tconv and green contours are gated as KLRG1-Tconv. (**C**) Quantitation of cytokine frequency across NPK-C1 and MCA205 tumors. Wilcoxon matched-pairs signed rank test was performed with Holm-Sidak multiple testing correction. Pooled data is plotted from two independent experiments; NPK-C1, *n* = 1-5 per replicate; MCA205 *n* = 4-5 per replicate. (**D**) Conceptual schematic of the T cell micro-suppression assay. (**E**) Representative examples of proliferation peaks of CTV-stained CD4+ T responder (Tresp) cells normalized to mode. Light grey peak represents peak 0 (non-dividing cells). (**F**) Quantitation of the cell division from each of the effector populations tested, represented as division index. Mann-Whitney test was used to assess statistical significance. Data shown are *n* = 7 that includes 2-3 technical replicates across 3 independent experiments. (**G**) Conceptual schematic of BMDC and T cell co-culture assay. (**H**) Quantitation of CD80+ and CD86+ BMDCs as fractions of total BMDCs (left) or XCR1+ BMDCs (right). Mann-Whitney test was used to assess statistical significance with Holm-Sidak multiple testing correction. Data shown are *n* = 7 including 2-3 technical replicates across 3 independent experiments. * = *p* ≤ 0.05, ** = *p* ≤ 0.01

### KLRG1+ Tconv are molecularly related to Tregs

To better understand the mechanisms underlying KLRG1+ Tconv function in tumors, we next employed RNA sequencing to interrogate the molecular wiring of tumor-infiltrating KLRG1+ Tconv relative to KLRG1-Tconv and Treg counterparts. For this, we established MCA205 tumors in *Foxp3*^RFP^ mice, harvested tumors at day 14 post-implantation, and flow sorted live CD45+CD3+CD4+ T cells split four ways based on expression of FoxP3 and/or KLRG1 (see Figure S3A for representative sorting strategy) then performed bulk RNA sequencing on each sorted population. We first compared the differential gene expression signature of KLRG1+ Tconv to KLRG1-Tconv and found KLRG1+ Tconv exhibited significant up regulation of genes corresponding to proteins detected in our prior flow-based assays including *Ctla4*, *Il10*, *Entpd1*, *Gzmb*, *Tnfrsf9*, *Ikzf2*, *Itgae*, *Havcr2*, and *Il2ra*, validating our prior findings and confirming the integrity of the dataset (Figure 4A). Of interest, *Foxp3* transcripts were detected in KLRG1+ Tconv despite no FoxP3 protein as measured via the red fluorescent protein (RFP) reporter used to sort the cells (Figure 4A; Figure S3A-B). Also supporting a connection between the KLRG1+ Tconv signature and Tregs, we observed upregulation of other Treg-related transcripts in KLRG1+ relative to KLRG1-Tconv such as *Areg*, *Ddit4*, *Tnfrsf18*, *Tnfrsf8* and *Ccr5.* We next compared KLRG1+ Tconv to KLRG1+ Tregs, and observed few differentially expressed genes, indicating high transcriptional similarity between these populations. Of the 6 significant DEGs, *Slamf7* and *Ifng* were upregulated in KLRG1+ Tconv versus KLRG1+ Tregs, supporting our prior *ex vivo* cytokine staining results and the cytotoxic signature characteristic of the KLRG1+ Tconv phenotype (Figure 4B). Interestingly, we also observed increased expression of *Il10* in KLRG1+ Tconv relative to KLRG1-Tregs (Figure 4C). In a global analysis, we found that KLRG1+ Tconv are more similar to Tregs than they are to KLRG1-Tconv, as shown by hierarchical clustering, likely driven by shared expression of core Treg genes including *Foxp3*, *Lrrc32*, *Il2ra*, *Ikzf2*, and *Itgae* (Figure 4D). The KLRG1+ Tconv and KLRG1+ Treg subsets shared a gene signature reminiscent of an effector tissue Treg cell state driven by *Ccl5*, *Il10*, *Ddit4*, *Ccr5*, *Areg*, *Ctla4*, and *Tigit*. Further supporting a putative shared identity between KLRG1+ Tconv and Tregs, gene set enrichment analysis (GSEA) comparing KLRG1+ Tconv to KLRG1-Tconv revealed genes sets related to IL-2/STAT5 signaling and response to various inflammatory cytokines enriched in KLRG1+ Tconv, and downregulation of Th1 cytokine production (Figure 4E). As KLRG1 expression has been associated with cellular senescence, we asked whether KLRG1+ Tconv were enriched in a senescence or senescence associated secretory phenotype (SASP) signature and found no significant enrichment in these cells relative to KLRG1-Tconv (Figure S3B). To more specifically ask whether KLRG1+ Tconv were enriched in a canonical Treg gene signature, we collated a 108-gene core Treg signature from published data of genes showing high correlation with FoxP3 in murine Tregs (*29*) and saw this signature was highly enriched in KLRG1+ Tconv versus KLRG1-Tconv (Figure 4E-F; FDR 1.15e-17). In support of this, we found KLRG1+ Tconv samples clustered closely to Tregs by PCA (Figure 4G). We also mapped expression of the core Treg genes to the four CD4+ sub-groups. We found that out of the 108 core Treg genes, the fewest genes with a z-score greater than 0.5 (averaged across the sub-groups) are found among the KLRG1-Tconv group, while the KLRG1+ Tconv cells have 35 overlapping genes,. In comparison, the KLRG1+ and KLRG1-Tregs have 47 and 54 overlapping genes, respectively (Figure 4H). Altogether, this data suggests that KLRG1+ Tconv are highly transcriptionally similar to Tregs, potentially indicating KLRG1+ Tconv are related to tumor-infiltrating Tregs.

**Figure 4:**
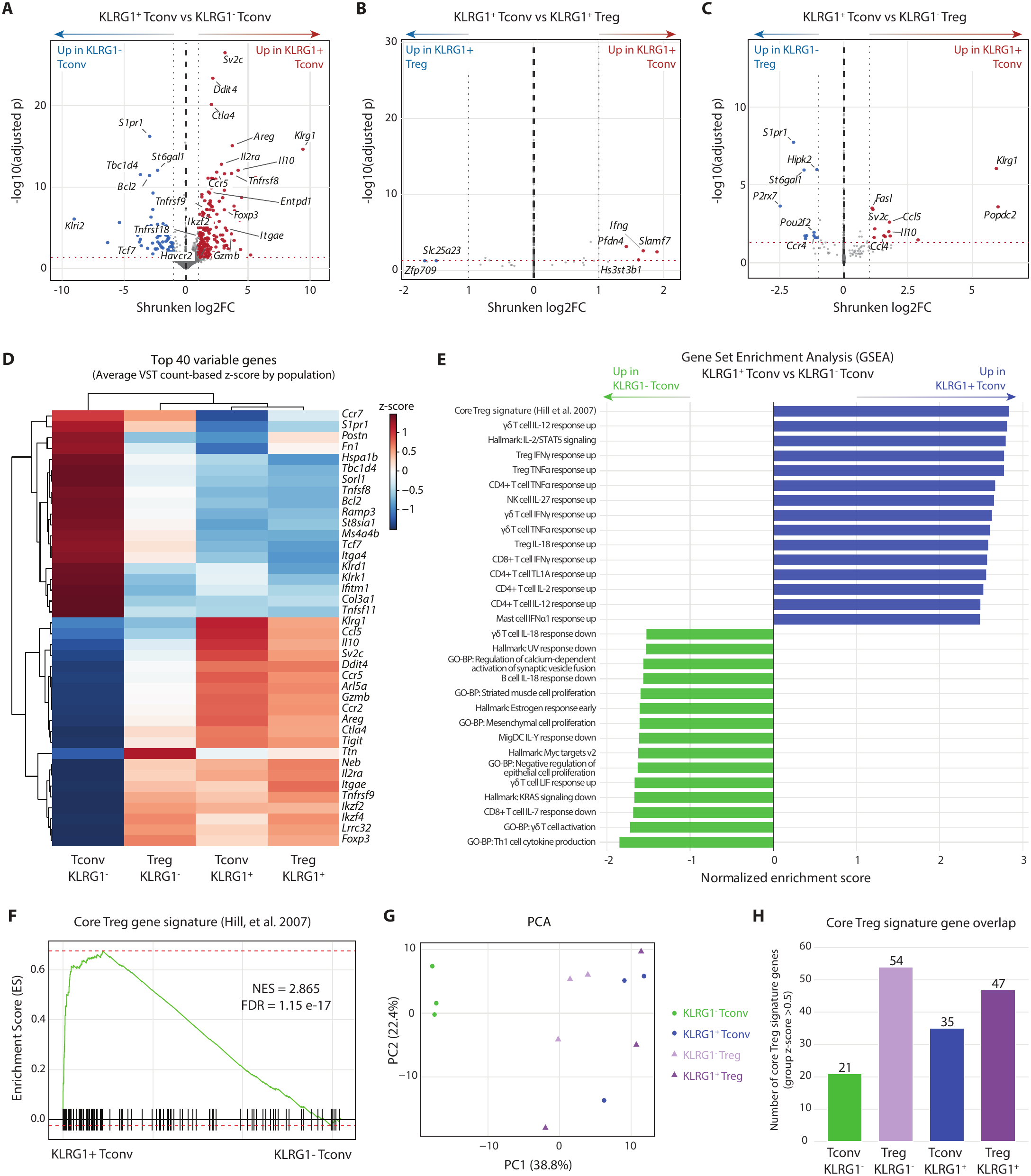
KLRG1+ Tconv are similar to Tregs at the molecular level. (**A**) Volcano plot of the differential expression analysis of KLRG1+ versus KLRG1-Tconv. (**B**) Volcano plot of the differential expression analysis of KLRG1+ Tconv versus KLRG1+ Treg. (**C**) Volcano plot of the differential expression analysis of KLRG1+ Tconv versus KLRG1-Treg. (**D**) Heatmap summary of the differential expression analysis. The top 40 variable genes across all groups are shown. Rows show the variance-stabilized transformation (VST) z-scored genes. (**E**) The top gene sets’ normalized enrichment score (NES) ordered by Benjamini-Hochberg adjusted-*p* value. The top 15 up and down regulated NES are shown based on the KLRG1+ versus KLRG1-Tconv DESeq2 comparison. Data shown used fgsea adaptive multilevel Monte Carlo procedure using the preranked Wald statistic from DESeq2. (**F**) Principal component analysis (PCA) of the VST gene counts for each biological replicate. (**G**) Running enrichment score curve for the core Treg gene signature collated from (*29*).(**H**) Gene overlap with the core Treg gene signature by group. Gene VST counts from the core Treg gene signature were averaged within each group and then the z-score for that gene in that group calculated across all groups. Genes with a z-score of >0.5 were considered having overlap with the gene set. An *n* = 3 biological replicates for sequencing was used. Log2 fold change of ≥1 in combination with an adjusted *p* ≤ 0.05 was the cutoff for statistical significance in differential expression analysis using DESeq2.

### KLRG1+ Tconv contain an IL-10 expressing, cytotoxic exTreg pool

Given the transcriptional similarity between KLRG1+ Tconv and Tregs despite lack of FoxP3 protein expression in KLRG1+ Tconv (as measured in our data by either flow cytometry antibody staining or a FoxP3-RFP reporter), we hypothesized that tumor infiltrating KLRG1+ Tconv are derived from Tregs but have lost FoxP3 expression; what have been called “exTregs.” To test this hypothesis, we employed an *in vivo* Treg fate-mapping approach by crossing the *Foxp3*^YFP-Cre^ mouse strain to the *Rosa26*^CAG-LSL-TdTomato^ (Ai14) strain (Figure 5A). In these animals, tdTomato reporter expression marks all cells that have expressed FoxP3, while YFP marks cells with active FoxP3 expression (Figure 5B). Therefore, CD4+ T cells with a YFP-tdTomato+ profile are considered exTregs. To determine whether KLRG1+ Tconv are more likely than KLRG1-Tconv to possess an exTreg phenotype, we subcutaneously implanted MCA205 tumors into *Foxp3*^YFP-Cre+/y^ *Rosa26*^CAG-LSL-TdTomato^ male mice, allowed tumors to grow to ∼200mm^3^, then harvested tumors and spleens and performed flow cytometry. After correcting for variable rates of background tdTomato signal known to be present in this strain (see Methods for detailed description and Figure S4A-B for example gating and quantitation), we observed a significant enrichment in exTregs (tdTomato+ and YFP-) within the KLRG1+ Tconv compartment versus KLRG1-Tconv (*p*=0.0002); approximately half of all KLRG1+ Tconv were exTregs (Figure 5C). Given that not all KLRG1+ Tconv were exTregs, we next sought to explore the functional capacities of KLRG1+ exTregs versus “never-Tregs” to better understand whether KLRG1+ exTregs were the subset of cells driving the cytotoxic and/or immunosuppressive phenotypes we previously observed. For this, we performed *ex vivo* re-stimulation and intracellular cytokine staining on MCA205-derived TILs from *Foxp3*^YFP-Cre^ *Rosa26*^CAG-LSL-TdTomato^ mice, leveraging an optimized fixation and permeabilization protocol that preserves both YFP and tdTomato signal and intracellular cytokines (*30*) (Figure S4C). We found that KLRG1+ exTregs expressed significantly more GzmB and IL-10 and reduced IFNγ as compared to KLRG1-exTregs. (Figure 5D-E). These findings suggest exTregs marked by KLRG1 are the exTreg fraction that maintains immunosuppressive functionality in this model. Next, we similarly compared KLRG1+ exTregs to KLRG1+ “never-Treg” Tconv cells. We observed a similar trend, where KLRG1+ exTregs are significantly enriched for expression of granzyme B (*p*=0.0078) and IL-10 (*p*=0.0391), while expressing significantly less IFNγ (*p*=0.0039) and TNFα (*p*=0.0078) as compared to KLRG1+ never-Tregs (Figure 5F and 5G). Together, these data suggest that the KLRG1+ Tconv compartment contains a large exTreg pool in MCA205 tumors, and that this KLRG1+ exTreg pool retains the GzmB+ IL10+ IFNγ/TNFα-low cytokine profile that may be responsible for the immunosuppressive function of KLRG1+ Tconv cells.

**Figure 5:**
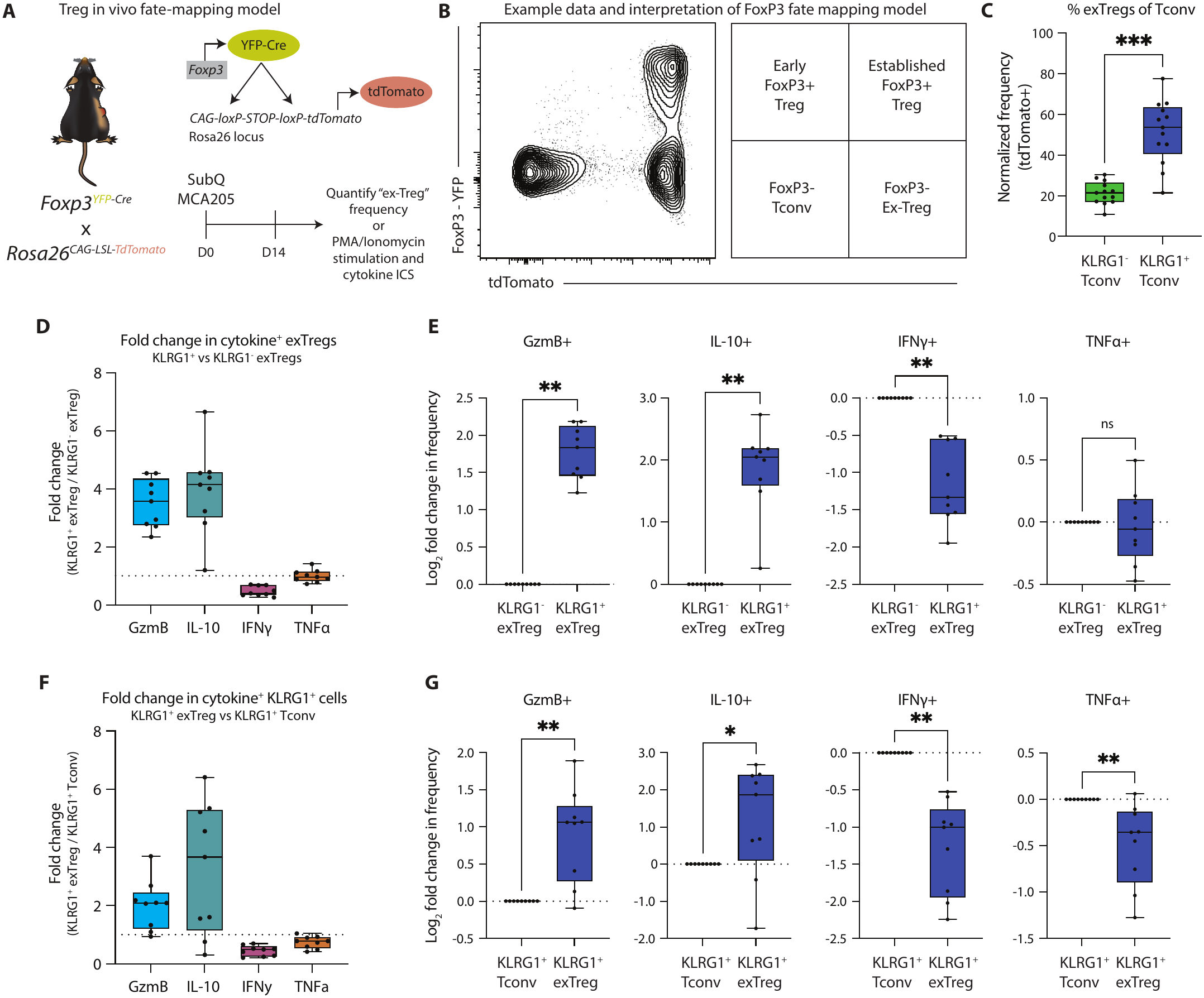
The KLRG1+ Tconv population contains a large IL-10+, GzmB+ exTreg pool. (**A**) Conceptual schematic of the fate mapping mouse system and timeline for fate-mapping experiments. (**B**) Example flow plot and interpretation for defining exTregs in the fate mapping system. (**C**) Quantitation of normalized tdTomato+ exTreg frequencies in KLRG1+ and KLRG1-Tconv cells. Frequencies were normalized on a per-mouse basis to the background frequency of tdTomato in CD8+ and CD3-cell populations (see Supplemental Figure 4A and 4B for more details). Wilcoxon matched pairs signed rank test was performed. Data shown include pooled data from groups of *n* = 6-7 mice from two independent experiments. (**D**) Summary plot of the fold change in cytokine-positive frequency of KLRG1+ exTregs to KLRG1-exTregs. The dashed line represents a log fold change of 1. (**E**) Quantitation of the log2-transformed fold change of each cytokine, normalized to the KLRG1-exTreg group. Wilcoxon matched pairs signed rank test was used for statistical analysis. Data shown include pooled data from two independent experiments with groups of *n* = 4-5 mice. (**F**) Summary plot of the fold change in cytokine-positive frequency of KLRG1+ exTregs to KLRG1+ Tconv cells that are not exTregs. The dashed line represents a log fold change of 1. (**G**) Quantitation of the log2-transformed fold change of each cytokine, normalized to the KLRG1+ Tconv group. Wilcoxon matched pairs signed rank test was used for statistical analysis. Data shown include pooled data from two independent experiments with groups of *n* = 4-5 mice. * = *p* ≤ 0.05, ** = *p* ≤ 0.01

### KLRG1-targeted immunotherapy slows tumor development and changes the TME

Current antibody therapeutics that target KLRG1 as a T cell inhibitory checkpoint are engineered to block the receptor. Based on our findings that intratumoral KLRG1+ cells are enriched in immunosuppressive CD4+ Tconv and CD4+ Treg populations (Figure S5A), along with the linked etiology of suppressive Tregs and KLRG1+ Tconv, we hypothesized that KLRG1-targeted antibodies with depletory function may be effective in driving antitumor immunity. To test this hypothesis, we produced monoclonal antibodies targeting KLRG1 with either Fc-silencing mutations that abrogate binding to murine FcgRs (STR; L240S/E241T/G242R “STR”) (*31, 32*)or Fc-enhancing mutations that enhance binding to FcgRs (DLE; S245D/S336L/I338E “DLE”) (*33, 34*) on the murine IgG2b backbone (Figure 6A). We validated antibody functionality and binding activity through competitive binding assays and enzyme-linked immunosorbent assays (ELISAs) (Figure S5B-C). We then compared the relative efficacy of these antibodies in the immunotherapy-responsive MC38 tumor model (Figure 6B). Survival of mice receiving αKLRG1-DLE versus those receiving αKLRG1-STR was significantly increased (*p*<0.0001; Figure 6C), paired with significantly delayed tumor growth compared to αKLRG1-STR or PBS-treated mice (*p*=0.000125 and *p*=0.000358, respectively; Figure 6D-E). These data support the notion that KLRG1-depleting immunotherapy is superior to KLRG1-blocking immunotherapy. To investigate the immune correlates of response to αKLRG1-DLE, we performed 43-color spectral flow on tumors four days after the last dose of immunotherapy. Semi-supervised clustering and dimensionality reduction delineated major tumor-infiltrating immune subsets (Figure 6F-G). We observed a significant reduction in tumor CD4+ T cell frequencies (*p*=0.0017; Figure 6H-J) in mice receiving αKLRG1-DLE, largely accounted for by the reduction in tumor Treg frequencies, consistent with expression of KLRG1 across 65-80% of Tregs in MC38 tumors (*p*=0.016; Figure S5C). Of note, despite known expression of KLRG1 in subsets of effector CD8+ and NK cells, we did not observe significant reductions in the NK or CD8+ T cell compartments (Figure 6J). To further investigate the impact of αKLRG1-DLE on the Tconv compartment, we sub-clustered the Tconv cells. The clustering solution broadly identified a meta cluster capturing a canonical naive/memory/effector trajectory and a highly differentiated, activated effector meta cluster defined by extensive checkpoint expression and, in part, KLRG1 (Figure 6K-L). We observed significant depletion of the KLRG1+ effector Tconv in animals receiving αKLRG1-DLE, paired with increases in central memory, proliferating effector memory, and highly differentiated/activated KLRG1-effector Tconv (Figure 6M-O). To validate these findings, we repeated this experiment in the MCA205 model, where we observed similar trends in high KLRG1 expression in tumor Tregs, trending tumor Treg depletion without reductions in CD8+ or NK cells frequencies, reductions in the KLRG1+ Tconv compartment, and increases in memory CD4+ Tconv phenotypes (Figure S5D-K). Together, these data suggest that KLRG1-targeted depletion may be superior to KLRG1 blockade at slowing tumor growth and improving survival in murine tumors; response to depletion-competent anti-KLRG1 antibodies associates with partial Treg depletion and a shift in the CD4+ Tconv compartment marked by increases in proliferative memory and highly activated KLRG1-effectors.

**Figure 6:**
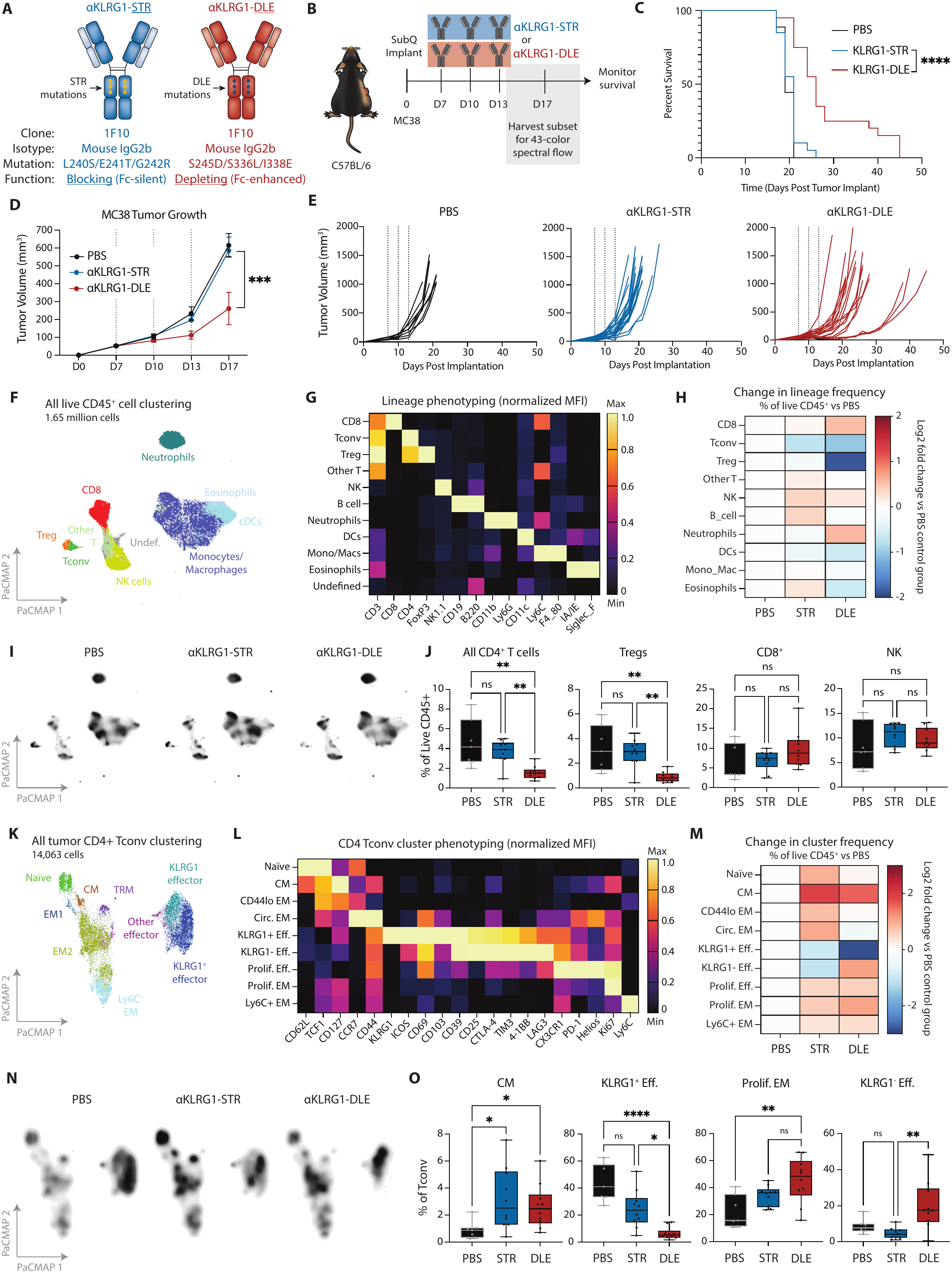
KLRG1-targeted depletion delays tumor growth and reduces tumor-infiltrating Treg frequency. (**A**) Graphical schematic of engineered anti-KLRG1 monoclonal antibodies (mAbs). (**B**) Schematic of *in vivo* therapeutic experimental timeline. (**C**) Kaplan-Meier curve showing survival of mice treated with therapeutics. Log rank test was performed for evaluating statistical significance. Pooled data from two independent experiments is shown. PBS group *n* = 3-5 per replicate; treatment groups *n* = 5-15 mice per replicate. (**D**) Average tumor growth of MC38-bearing mice in each respective treatment. Mann-Whitney test with Holm-Sidak multiple testing correction was used to assess statistical significance. (**E**) Individual mouse tumor growth curves from D. (**F**) PaCMAP of live CD45+ cells from treated MC38 tumors. Clusters are derived from FlowSOM. (**G**) Normalized heatmap of lineage-defining marker MFI across all live CD45+ cells. (**H**) Log2 fold change of each lineage cluster frequency in each treatment arm, normalized to the frequency in the PBS group. (**I**) Density plots of each treatment group overlaid on the lineage PaCMAP. (**J**) Frequency quantitation CD4+ T cells (left), Tregs (left-center), CD8+ T cells (right-center), and NK cells (right) as fractions of the live CD45+ clusters from each treatment group. Kruskal-Wallis one-way ANOVA with Dunn’s multiple testing correction was used for statistical testing. Data shown are pooled from two independent experiments. PBS control *n* = 3-4 per replicate; αKLRG1-STR and αKLRG1-DLE groups *n* = 5 per replicate. (**K**) PaCMAP plot of the CD4+ Tconv cluster from F. Clusters are derived from FlowSOM. (**L**) Normalized heatmap of phenotyping marker MFI in CD4+ Tconv across all treatment groups. (**M**) Log2 fold change of each CD4+ Tconv cluster frequency in each treatment arm, normalized to the frequency in the PBS group. (**N**) Density plots of each treatment group overlaid on the CD4+ Tconv PaCMAP. (**O**) Frequency quantitation central memory (CM; left), KLRG1+ effector (left-center), proliferating effector memory (Prolif. EM; right-center), and KLRG1-effector (right) cells as fractions of the CD4+ Tconv clusters from each treatment group. Kruskal-Wallis one-way ANOVA with Dunn’s multiple testing correction was used for statistical testing. Data shown are pooled from two independent experiments. PBS control *n* = 3-4 per replicate; αKLRG1-STR and αKLRG1-DLE groups *n* = 5 per replicate. ns = not significant, * = *p* ≤ 0.05, ** = *p* ≤ 0.01, *** = *p* ≤ 0.001, **** = *p* ≤ 0.0001

## DISCUSSION

CD4+ T cells possess a wide spectrum of functional roles in cancer ranging from productive help in priming CD8+ T cell anti-tumor responses, to active suppression of immune cells in the tumor microenvironment (*1, 5, 35*). In this study, we investigated the phenotypic and functional characteristics of CD4+ Tconv cells expressing the inhibitory receptor KLRG1, known primarily for its role in marking NK and cytotoxic CD8+ T cell states (*18, 36–38*). This phenotype was conserved across murine tumor models and in human ccRCC tumors and blood, where KLRG1 expression coincided with an activated, cytotoxic, and terminally differentiated phenotype in CD4+ Tconv. We found that KLRG1+ Tconv are functionally immunosuppressive and exhibit many transcriptomic, proteomic and functional features of FoxP3+ Tregs. Our *in vivo* lineage tracing studies demonstrated that the putatively immunosuppressive component of KLRG1+ Tconv was derived from Tregs that have lost FoxP3 protein expression but retain Treg-like functionality in tumors. Together, these findings challenge the paradigm that FoxP3 loss in Tregs causes loss of immunosuppressive function and provide an additional mechanism of intratumoral immunosuppression outside of canonical CD4+ FoxP3+ Tregs cells (*3, 6, 39–41*), and narrow the scope of markers that can assist in identifying and characterizing exTregs in mouse and human tumors.

The canonical definition of Treg instability is loss of FoxP3 expression concurrent with gain of inflammatory function, most commonly in inflammatory environments (*9, 15, 41–43*). Our fate-mapping experiments found that KLRG1+ Tconv were composed of up to 50% or more “unstable” or exTreg cells that had lost FoxP3. However, as a whole this compartment was functionally immunosuppressive in *ex vivo* assays. Furthermore, it appears the exTreg component of the KLRG1+ Tconv pool expressed the highest IL-10 and least IFNγ and TNFα. These data support the notion that FoxP3 loss is not sufficient for loss of immunosuppressive function. Intriguingly, maintenance of an IL-10+ IFNγ ^lo^ functional state in the FoxP3-exTreg pool was restricted to exTregs expressing KLRG1 in our model, implicating KLRG1 as a potential marker of immunosuppressive exTregs. Relatedly, a non-canonical type 1 regulatory (Tr1) cell subset has been defined as CD4+ Tconv lacking FoxP3 and CD25 but expressing immunosuppressive factors such as IL-10 and TGF-β. These cells are classically marked by expression of CD49b and LAG3 (*3, 44*) with other reports associating Tr1 cells with expression of PD-1, CTLA-4, ICOS, TIM3, TIGIT, CD226, CD73, CD39, GzmB, IFNγ, and Eomes, among others (*45–47*). We observed overlap of these identifying features in KLRG1+ Tconv, including IL-10, GzmB, CTLA-4, ICOS, TIM3, CD39, and LAG3 at the protein level, as well as *Tigit*, *Ccl5*, and *Ccr5* at the transcript level. However, a distinct downregulation of CD49b, IFNγ, Eomes, and lack of TGF-β at both the protein and transcript level would challenge the supposition that the KLRG1+ Tconv cells we describe are bona fide Tr1 cells. Moreover, there is evidence that Tr1 cells can exert immunosuppressive effects via cytotoxicity directed towards APCs such as cDC1s (*6, 48*). Sultan *et al.* nominated LILRB4, CCL5, and GzmB as defining markers for these specific cells. We found upregulation of LILRB4 and GzmB at the protein level and *Ccl5* at the transcriptional level in KLRG1+ Tconv, however we did not observe DC-directed cytotoxicity by tumor-derived KLRG1+ Tconv *ex vivo*. In prior studies, APC-directed Tr1 cytotoxicity in humans was MHC-I dependent (*48*) but in mice was MHC-II dependent (*6*), suggesting that mechanisms of Tr1 and Tr1-like cytotoxicity require further investigation.

Our study has immediate implications for development of KLRG1-targeting antibodies for cancer therapy. To our knowledge, we are the first to directly compare “Fc-enhanced” depletion-competent anti-KLRG1 antibodies versus “Fc-silenced” blocking-only anti-KLRG1 antibodies of the same clone and parent isotype. Our data is consistent with published studies that showed KLRG1 blockade alone is insufficient to slow tumor growth as single agent, generally requiring combination with PD-1/PD-L1 blockade to achieve significant growth and survival improvement (*23–25*). However, KLRG1-targeted depletion was effective as single agent in MC38 tumors, where it depleted both KLRG1+ Tconv and Tregs, importantly, without observable depletion of CD8+ effectors or NK cells. The seemingly selective activity at depleting KLRG1+ cells in the CD4+ compartment requires further investigation. It is notable that a depleting anti-KLRG1 antibody has undergone clinical development -- mAb208 from Abcuro Inc. -- for targeted depletion of malignant T and NK cell lymphomas marked by KLRG1 expression (*49*). Whether Treg depletion contributes to the efficacy of this agent, or whether this agent would have efficacy in solid tumors, remains unclear. Our group and others have explored other traditional forms of Treg depletion through mechanisms such as Fc-enhanced anti-CTLA-4-mediated depletion (*27, 50–52*). Whether Fc-enhanced KLRG1-targeted therapy would be redundant with Fc-enhanced CTLA-4 blockade or would possess superior or inferior activity remains to be tested. It is also notable that anti-4-1BB agonist therapy has been previously shown to induce highly cytotoxic KLRG1+ CD4 T cells with both effector and memory potential that play a key role in driving anti-4-1BB-mediated antitumor immunity (*53–57*). While our phenotyping data show 4-1BB expression is common on KLRG1+ Tconv, further investigation is required to determine if 4-1BB signaling is required or dispensable for the exTreg etiology or immunosuppressive function of the spontaneously arising KLRG1+ Tconv in our models. Overall, our study suggests depleting KLRG1-expressing cells may be a more robust therapeutic approach versus blocking KLRG1 signaling *in vivo*.

We acknowledge that we were limited in the number and type of human tumor samples to phenotype the KLRG1+ Tconv compartment, which constrained our statistical power and capacity to assess how generalizable the KLRG1+ Tconv phenotype is in human tumors. However, emerging studies from independent groups link KLRG1 expression to a cytotoxic CD4+ T cell program (*19, 20*), providing orthogonal evidence of this claim. Additional studies are warranted to assess the functional capacity of human tumor-derived KLRG1+ Tconv to determine if these cells are competent to produce IL-10. Technically, our fate mapping approach was challenged by varying rates of background tdTomato expression, which has been noted in other studies (*58*). While we corrected for this through mouse-specific background normalization, further studies in inducible systems may have utility in validating our findings. In addition, KLRG1+ Tconv were quite rare in the tumor microenvironment, limiting our capacity to do elaborate *ex vivo* functional assays required to assess the putative roles of IL-10, GzmB, TGF-β, or other cytokines in inhibition of T cell suppression by KLRG1+ Tconv. Additionally, more cells would be required for a systematic comparison of KLRG1+ Tconv, Tregs, and Tr1 cells via transcriptional or, ideally, epigenomic profiling to examine relationships between KLRG1+ Tconv and canonical Tr1 cells. Finally, it remains unclear where in the tumor KLRG1+ Tconv reside, thus future studies to interrogate the spatial coordinates of KLRG1+ Tconv in tumors is needed, particularly in investigating putative relationships between KLRG1+ Tconv and TREM2+ TAMs, which remains an exploratory finding requiring further study.

In conclusion, this study has defined immunosuppressive characteristics and cytotoxic potential of the KLRG1+ Tconv population, a subset we found to share features with Tr1 cells and to be in part derived from Tregs. These findings shed light on pathways for CD4+ plasticity in tumors and indicate the cytotoxic CD4+ compartment in tumors may in some contexts possess immunosuppressive function. Our preclinical therapeutic data suggest that KLRG1-targeted depleting antibodies may thus be a superior alternative to KLRG1 blockade, as this approach in murine models depleted KLRG1+ Tconv and Tregs without undue killing of CD8+ or NK cell effectors in tumors. These findings warrant further investigation and validation in human cohorts and provide new avenues to study the etiology and functionality of cytotoxic CD4+ T cells in cancer.

## MATERIALS AND METHODS

### Study design

The objective of this study was to understand the etiology of KLRG1+ Tconv and their functional characteristics that contribute to tumor immune responses. We used well-established murine tumor models to phenotype these cells and correlate them with other tumor immunophenotypes. This was followed by 51-color spectral immunophenotyping of healthy human PBMCs and patient ccRCC samples, which established in-human evidence of a late-stage, cytotoxic effector phenotype of KLRG1+ Tconv cells in tumors. Genetic mouse models for FoxP3 reporters were used to isolate CD4+ cells for *in vitro* functional characterization, for transcriptomic analysis, and for FoxP3 fate-mapping, resulting in clearer functional insights into exTregs in murine tumors. Our therapeutic approach was also aimed at comparing depleting versus blocking immunotherapy and showed the benefits of a depleting approach to tumor immunotherapy. Altogether, this study utilized spectral flow cytometry (protein level analysis) as the primary methodology for phenotyping and functional characterization of cells, validated by RNA sequencing and genetic models for fate-mapping (molecular level analysis).

### Mice

Wild-type C57/BL/6J, BALB/cJ, FVB/NJ, *Rosa26*^CAG-LSL-TdTomato^ (Ai14 or B6.Cg-*Gt(ROSA)^26Sortm14(CAG-tdTomato)Hze^*/J), and Thy1.1 (B6.PL-*Thy1^a^*/CyJ) mice were purchased from Jackson Laboratory and acclimated to the Mayo Clinic Arizona vivarium for at least a week prior to use. *Foxp3*^RFP^ (FIR or C57BL/6-*Foxp3^tm1Flv^*/J) and RAG2-knockout (B6.Cg-*Rag2^tm1.1Cgn^*/J) mice were a kind gift from Dr. Khashayarsha Khazaie. *Foxp3*^YFP-Cre^ mice were a kind gift from Dr. Fotini Gounari. The *Foxp3*^YFP-Cre^ x *Rosa26*^CAG-LSL-TdTomato^ fate mapping mice were crossed in-house and hetero/homozygous tdTomato alleles and hemi/homozygous Cre alleles were deemed appropriate for experimental fate mapping. Mice beyond the F3 generation of inbreeding were not used. Male and female mice were used approximately equally across experiments, except for FVB/NJ mice which were exclusively male to model prostate MycCaP tumors. Mice were housed in accordance with the NIH Guide for the Care and Use of Laboratory Animals and the Association for Assessment and Accreditation of Laboratory Animal Care (AAALAC) International standards and regulations, holding at a maximum of 5 animals per cage, food and water provided *ad libitum*, with a 14-h light cycle. All animals were approximately 6-12 weeks old at time of use. All experiments and procedures described in this study were approved by the Mayo Clinic Institutional Animal Care and Use Committee (IACUC).

### Healthy human PBMCs

Anonymous healthy donor PBMCs were obtained from apheresis cones (*59*) upon request by the Division of Transfusion Medicine at Mayo Clinic Rochester. Briefly, cone volumes were diluted 1:1 with sterile, room temperature PBS prior to density centrifugation with Ficoll-Paque (1.077g/mL; Cytiva) for 30 minutes at 18°C with the brake set to 2. Cells were washed twice with cold PBS and counted. Aliquots of 2-50 x 10^6^ cells were cryopreserved in CryoStor CS10 cryopreservation medium (Sigma Aldrich). For us in flow cytometry, cryopreserved cells were thawed quickly in a 37°C water bath, added to 10mL of warm R10 media, and pelleted at 300xg, 20°C, for 5 minutes before counting. Up to 2 x 10^6^ live cells were used in the spectral flow staining protocol for samples or single-color controls.

### Human ccRCC tissue

Patients undergoing radical or partial nephrectomies as part of standard care for ccRCC were consented to IRB #08-000980 at Mayo Clinic Arizona. Tissue was received from Surgical Pathology and processed in the same manner as described under the spectral flow cytometry method section below. Single-cell suspensions were counted after digestion and up to 5 x 10^6^ live cells were stained for spectral flow cytometry.

### Tumor cell lines

B16-F10 cells were a kind gift from Dr. Jessica Lancaster. Renca cells were purchased from ATCC. MC38 was purchased from Kerafast. MB49 and MCA205 were purchased from Millipore Sigma. MycCaP cells were a kind gift from Dr. Charles Drake. NPK-C1 cells were a kind gift from Dr. Cory-Abate Shen. All cell lines were verified as free of pathogen and cell contamination free through IDEXX BioAnalytics. B16-F10, MC38, and MB49 cells were grown in Dulbecco’s modified MEM (DMEM) with 10% FBS, 100U/mL penicillin, and 100mg /mL streptomycin. RENCA, MycCaP, and NPK-C1 cells were grown in RPMI-1640 with 10% FBS, 100U/mL penicillin, and 100mg/mL streptomycin. MCA205 cells were grown in RPMI-1640 with 10% FBS, 100U/mL penicillin, 100mg/mL streptomycin, 1X MEM non-essential amino acids (Corning) and 55mM b-mercaptoethanol.

### Animal studies

For tumor implantation, cells in culture at ∼70-90% confluency were harvested with 0.05% trypsin-EDTA, washed with cold PBS, counted and resuspended at 10 x 10^6^ cells/mL in cold PBS. On day 0, mice were implanted subcutaneously in the right flank with 1 x 10^6^ cells in 100mL or 0.5 x 10^6^ cells in 50mL in the case of B16-F10. Tumors were measured with digital calipers in X, Y, and Z dimensions to calculate tumor volume every 2-3 days once tumors became palpable. For tumor model immune profiling, tissue was harvested once group averages were at least 150 mm^3^. For experiments where cells were sorted, tumor volumes were at least 200mm^3^, with 2-3 mice used to source cells that were pooled post-sorting for *in vitro* experiments or maintained separate for bulk RNA sequencing. For therapeutic experiments, tissue was harvested 4 days after the last dose of immunotherapy for immune profiling with groups of mice that were randomly pre-determined at the outset of tumor implantation. Remaining mice were used for following therapeutic survival. Mice were deemed eligible for euthanasia when tumor volume surpassed 1000mm^3^ or tumor ulceration exceeded 5mm in diameter. Independent replicate experiments were performed with cohort sizes of n = 3-5 for immune profiling by flow cytometry and n = 5-15 for monitoring tumor growth and survival in response. Data from all mice are included in flow and survival analyses, other than objective outliers (mice where <10,000 live immune cells were acquired for immune profiling by flow cytometry or PBS-treated mice had abnormal tumor growth, meaning tumor did not become palpable or tumor growth was slowed beyond typical for that model; n = 2).

### Spectral flow cytometry

#### Tissue processing

Following euthanasia, tumor mass was recorded and up to 50mg of tumor tissue was diced and dissociated in X-VIVO 15 media (Lonza) supplemented with 0.16mg/mL of DNaseI (Millipore Sigma) and 1mg/mL of collagenase D (Millipore Sigma) with rocking at 37°C for 30 minutes. Digestion reactions were quenched by adding 5mL of R10 media and filtered through 70μm filters. Single-cell suspensions of dissociated tissue were then centrifuged at 300xg for 5 minutes at 4°C and resuspended in 200uL of cold R10 and transferred to 96-well, round-bottom plates for spectral flow cytometry staining.

#### Spectral staining protocol

Cells were washed twice with PBS. Dead cells were stained with 100 μL + Zombie NIR (BioLegend) or Live/Dead Fixable Blue dyes (1:1000 in PBS, Invitrogen). Unless otherwise stated, this and all other staining steps were performed for 30 minutes at 20°C with gentle shaking at 400rpm on a Thermomixer. Samples were washed twice with FACS buffer (PBS + 3% FBS + 1mM EDTA + 10mM HEPES; all reagents from Corning) then resuspended in FACS with either TruStain FcX or TruStain FcX PLUS (BioLegend) at 1:50. Surface antibodies were prepared in a master mix at optimal dilutions in FACS supplemented with Brilliant Stain Buffer Plus (BSB Plus; 10 μL per sample; BD Biosciences) and True-Stain Monocyte Blocker (1:20; BioLegend) and added on top of the samples in blocking solution. After surface staining, samples were washed twice with FACS buffer and fixed with 100μL of FoxP3 Fixation/Permeabilization Kit buffer (ThermoFisher Scientific). Samples were then washed twice with 1X permeabilization buffer (1X PW) and resuspended in 1X PW. Intracellular antibodies were prepared in a master mix at optimal dilutions in 1X PW with BSB Plus and True-Stain Monocyte Blocker and added on top of the resuspended samples. Plates were sealed and intracellular staining proceeded overnight at 4°C on an orbital shaker. The next day, samples were washed twice with FACS buffer, resuspended in 200μL FACS buffer and immediately acquired on a Cytek Aurora 5-laser (UV/V/B/YG/R) instrument. Splenocytes or PBMCs and SpectraComp (Slingshot Bio) or UltraComp (Invitrogen) beads were stained in parallel with samples and used as single color controls for unmixing.

#### Analysis of flow cytometry data

Analysis was performed using FlowJo and the R computational environment, version 4.4.3. Live, single cell, CD45+ events were manually gated using standard gating practices (see Supplemental S1A and S2A for representative plots from mouse and human tissues, respectively). For high dimensional analysis, samples with fewer than 10,000 live CD45+ events were excluded from downstream analysis. Otherwise, up to 50,000 live CD45+ events were down sampled and concatenated. Immune lineages were defined and annotated in a semi-supervised manner using FlowSOM, over-clustering based on lineage-defining markers and then merging based on lineage markers, as described previously(*27*). Clusters were then embedded into two-dimensional space using the same lineage-defining markers and the PaCMAP (*60*) plugin in FlowJo. An additional round of sub-clustering on immune lineages was then performed in a similar manner using phenotypic markers before embedding into two-dimensional PaCMAP space for visualization. Cell numbers/frequencies and marker MFI were exported for clusters and other defined groups based on keywords for statistical analysis and further visualization. Data was scaled with min-max normalization or using arcsinh transformation with a cofactor of 1500. For non-high dimensional flow experiments, gated population frequencies were exported and plotted as a proportion of a parent population (see figures and figure legends for details) or as fold change in frequencies between populations of interest.

### Ex vivo cytokine stimulation

#### Tissue processing

For *ex vivo* cytokine stimulations, NPK-C1 tumors were harvested at endpoint or MCA205 tumors were harvested at day 14 post-implantation and processed as described for spectral flow cytometry, with some modifications. Whole tumors were digested before mononuclear cell enrichment via density centrifugation using Ficoll-Paque PREMIUM 1.084 (Cytiva) for 20 minutes at 18°C with the brake set to 2. The mononuclear cell fraction was then washed, resuspended with R10, and counted.

#### PMA/Ionomycin stimulation and staining

Up to 2 x 10^6^ enriched mononuclear cells were then aliquoted into a 96-well, round bottom plate. A PMA/ionomycin solution (100ng/mL and 1μg/mL for final concentrations, respectively) were added on top of the cells and incubated for 30 minutes at 37°C with 5% CO_2_. Brefeldin A and Monensin (5μg/mL and 2μM, respectively; BioLegend) were added on top of the stimulated cells and incubated a further 3.5 hours. Samples were then washed once with PBS, and dead cells were stained with 100μL + Live/Dead Fixable Blue dye (1:1000 in PBS). Samples were washed twice with FACS buffer and then fixed with 100μL of FoxP3 Fixation/Permeabilization Kit buffer. All further staining steps follow the spectral staining protocol method. Unstimulated cells or fluorescence minus one (FMO) controls were used to determine positivity for cytokines. For *ex vivo* cytokine stimulation from fate-mapping tumor, see fate-mapping methods.

### Suppression assay

#### Cell sorting

Whole MCA205 tumors were harvest at day 14 (∼200mm^3^) and processed to a single-cell suspension as described for flow cytometry staining. Filtered cells were resuspended in 1mL of FACS buffer in a 5mL polypropylene tube and surface stained for 30 minutes on ice with occasional mixing (see Flow Panels spreadsheet for panel and dilutions). Cells were washed twice with FACS buffer and resuspended in 2-4mL of FACS buffer for cell sorting. Helix Green viability dye (1:10,000 in FACS buffer; BioLegend) was added immediately before beginning sorting. Sorting was performed on a Sony MA9000 instrument to isolate CD4+ KLRG1+ or KLRG1-Tconv or Tregs. Mouse splenocytes were processed simultaneously and used for single-color controls, and for gating strategy design.

#### Micro-suppression assay

Splenocytes from RAG2-KO mice and Thy1.1 mice were isolated by mechanical dissociation of spleens and passing through 70μm filters. Single cell suspensions were ACK lysed with 1mL of ACK buffer (Quality Biological) and resuspended in 0.5mL of R10 media. Thy1.1 splenocytes were then enriched for naïve CD4+ T cells using the MojoSort Mouse CD4 Naïve T cell isolation kit (BioLegend) and stained with CellTrace Violet (CTV; Invitrogen). Final cell suspensions (including FACS sorted tumor-infiltrating CD4+ T cell populations) in R10 were counted and up to 5000 tumor-infiltrating CD4+ effectors (effectors) were mixed at the following ratios before serial diluting while maintaining consistent CTV-labeled responders and antigen-presenting cells (APCs) from RAG2-KO mice: 1:1 for responders and APCs, with one suppressing effector for every two responder cells (1:2). Functional anti-CD3 antibody (BioLegend) was added at a final concentration of 1μg/mL and mouse IL-2 (Peprotech) was added to a final concentration of 5ng/mL. Cells were incubated ∼86 hours at 37°C with 5% CO_2_ before a surface stain for 30 minutes on ice (see Flow Panels spreadsheet for details). Cells were then washed once with FACS buffer, resuspended in 200μL of FACS buffer and immediately acquired on a Cytek Aurora instrument. CTV-labeled responders without anti-CD3 were used to determine a non-proliferative population control. Proliferation of live, CTV-stained Thy1.1+ cells were analyzed in FlowJo using the proliferation modeling tool and division index (calculated as total # of divisions / # of cells at start of culture) used as the main output for proliferation. Suppression effects were seen diluted out as low as the 1:4 dilution of effectors.

### Bone marrow-derived dendritic cell (BMDC) co-culture assay

#### Dendritic cell differentiation from bone marrow

Bone marrow from WT C57/BL/6J mice was extracted from femur and tibia bones by cutting one end and centrifugation for 5000xg for 5 minutes at 4°C in a 0.5mL tube containing an 18-gauge needle hole in the bottom nested within a 1.5mL tube. Extracted bone marrow was ACK lysed and resuspended in 1mL of RPMI-1640 containing 10% FBS, 100U/mL penicillin, 100μg/mL streptomycin, 25mM HEPES, 1X MEM non-essential amino acids (Corning), and 55μM β-mercaptoethanol. Cells were counted and plated at 6.25 x 10^6^ cells/mL with 800ng/mL of Flt3-ligand-Fc (BioXCell) and 200ng/mL of SCF (Peprotech) in 10cm dishes. Cells were incubated at 37°C with 5% CO_2_ for 3 days. At day 3, media was removed, replaced containing only 800ng/mL Flt3-ligand-Fc, and cells incubated a further 7 days. Non-adherent and loosely adherent cells were harvested by gentle washing with medium without Flt3-ligand-Fc at day 10 for downstream experiments.

#### BMDC and T cell Co-culture

MCA205 tumors were harvest and CD4+ T cell populations sorted as described for suppression assays. BMDCs from day 10 were then mixed 1:1 with sorted KLRG1+/-Tconv or Treg cells and incubated overnight at 37°C with 5% CO_2_ in RPMI-1640 containing 10% FBS, 100U/mL penicillin, 100μg/mL streptomycin, 25mM HEPES, 1X MEM non-essential amino acids, 55μM β-mercaptoethanol, plus 5ng/mL of mouse IL-2. The next day, cells were surface stained for 30 minutes on ice (see Flow Panels spreadsheet for details). Cells were then washed once with FACS buffer, resuspended in 200μL of FACS buffer and immediately acquired on a Cytek Aurora instrument. Inactivated BMDCs alone without T cells were used as a baseline for CD80 and CD86 expression. For experiments explicitly evaluating BMDC cytotoxicity (Supplemental figure 2), BMDCs were pulsed over night with equal volumes of either PBS or MCA205 tumor lysate + 40ng/mL of murine IFNγ (Peprotech) and 100ng/mL of LPS (*E. coli* O55:B5 from Fisher Scientific) to induce activation and antigen presentation overnight before co-culturing the next day with tumor-derived CD4+ T cell populations as described in the cell sorting methods. MCA205 tumor lysate was generated by repeated freeze/thaw cycles of MCA205 tumor cells at 10 x 10^6^ cells/mL until viability was less than 5% as assessed by Trypan Blue exclusion. Approximately 3 tumor cells worth of lysate was added for every BMDC for the overnight incubation. Cytotoxicity assessment was performed independently two times while the CD80/CD86 quantitation was performed 3 times.

### Monoclonal antibody production and purification

The 1F10 anti-KLRG1 variable region sequences were kindly shared by Dr. Laurent Brossay and Dr. Eric Vivier/Innate Pharma. Briefly, the variable regions for the heavy chain (HC) and light chain (LC) were added to the constant regions for a mouse IgG2b HC and a mouse kappa LC, taken from the IMGT database and based on allele 01. The L240S/E241T/G242R and S245D/S336L/I338E mutations for silencing or enhancing Fc receptor binding, respectively, were introduced at the nucleotide level. The Kozak consensus sequences and secretion signal peptide were added upstream of the variable regions before codon optimization for Chinese hamster ovary (CHO) cells. Final gene constructs were synthesized by Twist Bioscience and cloned into the pTwist-CMV-BG-WPRE-Neo expression plasmid. Plasmids were propagated in *Escherichia coli* (DH5α strain, New England Biolabs) using carbenicillin at 50 μg/mL and purified using the GenElute™ HP Endotoxin-Free Plasmid Maxiprep Kit (Sigma-Aldrich). The ExpiCHO Expression system (Gibco) was used to produce the monoclonal antibodies, according to the manufacturer’s “Max Titer Protocol”. In short, ExpiCHO-S cells were grown in defined ExpiCHO Expression Medium into log phase growth (∼7 – 10 x 10^6^ cells/mL), diluted with medium to ∼6 x 10^6^ cells/mL, and transfected with 0.8-1μg of total plasmid DNA (HC to LC ratio was ∼1:1) using ExpiFectamine CHO Reagent in OptiPRO serum-free medium. Post-transfection cells were incubated ∼18-24 hours at 36.5°C with 8% CO_2_ and 120 rpm shaking before adding the ExpiCHO Enhancer and Feed and a shift to 32°C and 5% CO_2_ for the remaining 14-day expression run. A second feed was added at day 5 post-transfection.

#### Monoclonal antibody purification

ExpiCHO cell supernatant was harvested at day 14 post-transfection for monoclonal antibody purification. Cell debris was pelleted by centrifugation for 30 minutes in 4°C at 5000xg. The clarified supernatant was filtered through 0.45 μm vacuum filters before capturing monoclonal antibodies with MabSelect Protein A affinity resin (Cytiva) in a gravity flow system. Unbound molecules were washed out with at least 10X column volume (CV) with PBS pH 7.4 before eluting the monoclonal antibodies with 50 mM sodium citrate, pH 3.0. The eluate was quickly neutralized to pH 7 with a 20% volume addition of 1 M Tris-HCl, pH 8.0. The purified monoclonal antibodies were then concentrated and buffer exchanged into PBS pH 7.4 using 100 kDa MWCO centrifugal filters (MilliporeSigma). Concentrated monoclonal antibodies were then filter-sterilized with 0.2 μm syringe filters before aliquoting and freezing in -80°C until used in downstream experiments. Filter-sterilized antibodies were pathogen tested via IDEXX Bioanalytics as described for tumor cell lines.

#### Monoclonal antibody quantitation and functional validation

A frozen aliquot of antibody was thawed and concentration estimated with A_280_ on a Nanodrop before further quantitation via sandwich ELISA, detecting a fully assembled IgG molecule using a strategy previously implemented to quantify mAb concentration (*61, 62*) using anti-mouse commercially available mouse IgG2b control as a standard (Genscript). Concentration and purity were also validated via SDS-PAGE after ELISA quantitation. To validate antibody target binding, a competitive binding assay on mouse splenocytes was performed. Briefly, a mouse spleen was mechanically dissociated, passed through a 70 μm filter, red-blood cells lysed, and ∼1 x 10^6^ cells aliquoted into 5 mL polypropylene FACS tubes. Cells were then incubated with saturating concentrations (10 μg/mL) of the house-made anti-KLRG1-STR/DLE antibodies, the control anti-KLRG1 2F1 antibody (BioXCell), or PBS and incubated on ice for 30 minutes. Then the master mix containing lineage markers (CD45, CD3, NK1.1, KLRG1, etc. see Flow Panels spreadsheet for details) was adding on top and further incubated 30 minutes. Antibodies were then washed with 2X washes with 1 mL of FACS buffer, followed by centrifugation of 5 minutes, 300xg, at room temperature. Cells were resuspended in 200 μL of FACS buffer, data acquired on the Cytek Aurora, and live CD3+ KLRG1+ and NK1.1+ KLRG1+ population frequencies calculated. To validate the STR and DLE mutations modulate binding to mouse CD16.2 (FcγRIV; R&D Systems), a sandwich ELISA assay was performed where purified mouse CD16.2 (was coated at 2μg/mL overnight on a high-binding polystyrene plate in carbonate-bicarbonate buffer, pH 9.6. Then either commercially available WT mouse IgG2b, anti-KLRG1 STR/DLE were serially diluted and incubated. Mouse IgG2b binding to CD16.2 was detected with goat anti-mouse kappa conjugated to horseradish peroxidase (HRP; Southern Biotech) at 0.5 μg/mL. TMB substrate and stop solution (50 μL at each step; BioLegend) was used to develop the plate and halt the enzymatic reaction. The absorbance at 450 nm was read using a plate reader immediately. All ELISA steps used Tris-buffered saline with 0.05% Tween-20 (TBST; Cell Signaling Technology) as the washing agent, performed twice and between each incubation with 200 μL per well. ELISA blocking steps were performed with 200μL per well of 5% dry milk (ThermoFisher) dissolved in TBST. ELISA incubation steps other than the initial plate coating were performed at 37°C for 1 hour with 50 μL per well.

#### Antibody treatments

For *in vivo* antibody treatment, filter sterilized, in-house made αKLRG1-STR or αKLRG1-DLE (both recombinantly-produced mouse IgG2b) were thawed on ice, diluted in cold, sterile PBS, and administered to mice via intraperitoneal (IP) injection to tumor-bearing mice. Each mouse received 100μg of the designated antibody at days 7, 10, and 13 post-tumor implantation.

### Bulk RNA sequencing and analysis

#### RNA Isolation and library preparation

For bulk RNA sequencing, MCA205 tumor-infiltrating CD4+ KLRG1+/- Tconv or Tregs were FACS sorted as described for the suppression and BMDC co-culture assays. Cells were washed twice with sterile PBS, QIAzol lysis reagent (QIAGEN) added to the cells, and frozen until ready for RNA extraction. RNA extraction was performed using the miRNeasy Micro Kit (QIAGEN) according to manufacturer instructions. Final extracted RNA concentration was estimated on a Nanodrop and ∼90ng of total RNA was used as input for library preparation. The Stranded mRNA Prep Kit (Illumina) was used for library preparation according to manufacturer’s instructions. Final library concentrations were quantified with a Qubit fluorometer, sizes estimated by Tapestation analysis, and libraries normalized to a pooled concentration of ∼3nM. Approximately 40-50 million reads per sample were obtained on a NovaSeq instrument at Azenta Life Sciences.

#### Differential expression analysis

Raw fastq files were uploaded to galaxy-main.usegalaxy.org and paired end reads aligned to the NCBI RefSeq GCF_000001635.27-RS_2024_02 mm39 genome assembly with the associated gtf using RNA STAR (*63*) and default parameters. Stranded gene counts (--quantMode GeneCounts) from RNA STAR were then exported and imported into R (version 4.4.3) and filtered to retain protein-coding, non-mitochondrial genes using biotype annotations from EnsDb.Mmusculus.v79 that contained at least 20 counts across the 3 replicates used for each cell population. An initial QC comparison between groups revealed several myeloid and tumor genes (*Apoe, Lyz2, Col3a1,* etc.) that were skewing the analysis in non-biological direction (sorting purity for some CD4+ T cell populations was ∼85% prior to RNA extraction). To help reveal true biology, not contaminating counts, 32 genes (25 after the low count filtering) defining tumor-associated macrophages and other myeloid populations (*64*) were used to generate a myeloid covariate where each gene was z-scored across all samples, averaged in each sample, and standardized to a mean of zero and standard deviation of one before DESeq2 modeling. DESeq2 1.46.0 (*65*) was then used to model the raw counts using standard size-factor normalization, dispersion estimation and negative-binomial modeling with a design of “∼ myeloid covariate + group” where group was KLRG1+/- Tconv or Treg (four groups total). Volcano plots were generated using the variance-stabilized counts in the DESeq2 model using Wald tests with Benjamini-Hochberg adjustments for differential expression p-values and fold changes shrunk using apeglm. The top variable gene heat map was generated using pheatmap and the group mean averages, standardized across the four group means, and hierarchically clustered using Euclidean distance and complete linkage applied to the z-scored gene matrix.

#### Gene set enrichment analysis

Pre-ranked GSEA was performed on the KLRG1+ Tconv versus KLRG1-Tconv comparison in the DESeq2 model using fgsea (*66*) 1.32.2 and msigdbr 26.1.0 in R (version 4.4.3) and gene sets in mouse MSigDB 2026.1.Mm. Gene sets needed a minimum of 5 and a maximum of 500 genes for our analysis.1,227 gene sets were tested spanning all mouse Hallmark(*67*), all M7 immunologic signatures(*68*), all M5 mouse phenotype tumor ontology sets, and selected M5 gene ontology biological processes filtered based on the following terms: CD4, CD8, Treg, regulatory T cell, CD25, T helper, alpha beta, memory, effector, cell death, apoptosis, activation, and proliferation. Additionally, a manually collated core Treg gene set based on genes that correlate with FoxP3 expression (≥ 0.3) and map to a Treg signature (fold change ≥ 2.0) based on the microarray supplement from (*29*) resulted in an additional gene set called “Core Treg signature” with 108 genes. Signed Wald statistics were used as the ranking metric, Benjamini-Hochberg multiple-testing correction applied and an FDR of < 0.05 set as the significance threshold. Of 388 significant sets, the top 15 positive and negative NES sets were plotted based on significance. For gene overlap with the core Treg gene signature, VST counts for each core Treg gene was averaged across samples in each group before calculating a z-score across the four groups for each gene. Genes with z ≥ 0.5 were considered strong candidates for having “overlap” with core Treg genes. A supplementary senescent signature GSEA analysis with the same analysis parameters was performed using the following gene sets: SAUL_SEN_MAYO(*69*); select M2 CP:Reactome sets including: DNA_DAMAGE_TELOMERE_STRESS_INDUCED_SENESCENCE, CELLULAR_SENESCENCE, OXIDATIVE_STRESS_INDUCED_SENESCENC, and SASP; other M5 gene ontology biological processes sets: POSITIVE_REGULATION_OF_CELLULAR_SENESCENCE, NEGATIVE_REGULATION_OF_CELLULAR_SENESCENCE, and REPLICATIVE_SENESCENCE.

### FoxP3 fate-mapping

#### ExTreg quantitation

MCA205 tumors were implanted and tissue processed as described above. For exTreg quantitation, filtered single-cell suspensions from tissues (spleens and tumors) were resuspended in 1mL of FACS buffer and surface markers stained for 30 minutes on ice with occasional mixing (see Flow Panels spreadsheet for more details). Cells were washed twice with FACS buffer, resuspended in 200-500μL and acquired immediately on a Cytek Aurora instrument. Approximately 1 x 10^6^ events were acquired for splenic tissue and whole tumors were acquired. Live, CD45+ events were down sampled to 100,000 events for final data analysis. Background tdTomato frequencies in non-Cre expressing cell types (CD3+ CD8+ and CD3-cells) were quantified in spleen and tumor tissue and used to normalize CD4+ tdTomato frequency by subtracting the average off-target frequency from the CD4+ frequency on a per-mouse basis.

#### Fate-mapping ex vivo cytokine stimulation

For *ex vivo* cytokine stimulation in the fate mapping system, MCA205 tumors were processed as described in the *ex vivo* cytokine methods, with a few modifications. Whole tumor, single-cell suspensions were stimulated in 6-well plates, with the same 30-minute lead-in PMA/ionomycin stimulation followed by 3.5 hours of Brefeldin A and Monensin secretion inhibition. After stimulating, cells were washed twice with PBS and cells split into multiple wells to account for beginning tumor mass (∼50mg of tissue per well for staining). Dead cells were stained with 100μL of Zombie NIR (1:1000 in PBS). To preserve the fluorescent protein signals while simultaneously staining for cytokines, the dish soap protocol (*30*) was implemented as an alternative fixation and permeabilization method. Briefly, cells were fixed with 100μL of fixation buffer (2% paraformaldehyde, 0.05% Dawn dish soap, and 0.5% Tween-20) for 30 minutes at 20°C with gentle shaking at 400rpm on a Thermomixer. Cells were then washed twice with permeabilization buffer (0.05% Dawn dish soap in PBS; here 1X DPW) and resuspended in 1X DWP. Intracellular antibodies were prepared in a master mix at optimal dilutions in 1X DWP with BSB Plus and True-Stain Monocyte Blocker and added on top of the resuspended samples. Plates were sealed and intracellular staining proceeded overnight at 4°C on an orbital shaker. The next day, samples were washed twice with FACS buffer, resuspended in 200μL FACS buffer and immediately acquired on a Cytek Aurora 5-laser (UV/V/B/YG/R) instrument. Splenocytes and SpectraComp beads were stained in parallel with samples and used as single color controls for unmixing. FMO controls were used to define cytokine positive cells during data analysis.

### Statistical analysis

All tests on flow cytometry data for population frequencies, targeted correlation plots, and tumor growth and survival curves were performed in GraphPad Prism 11.1.0. Correlation matrix analysis, using Spearman’s correlations, and RNA sequencing analysis was performed in the R computational environment (version 4.4.3). Where appropriate, Kruskal-Wallis, Mann-Whitney, Wilcoxon matched pairs signed rank, or two-sided paired t tests were performed. When appropriate, Holm-Sidak, Benjamini-Hochberg, or Dunn’s multiple testing correction was applied. Specific tests and population sizes are detailed in the figure legends. Statistical significance was defined as *p* values or false discovery rate (FDR) of less than 0.05. Bars in box and whisker plots represent the range of the data while the box represents the 25-75^th^ percentile range and the solid line represents median. Unless stated otherwise, *p* values are indicated as: ns = not significant, * = *p* ≤ 0.05, ** = *p* ≤ 0.01, *** = *p* ≤ 0.001, **** = *p* ≤ 0.0001.

## Supporting information

Supplemental Tables - Flow Cytometry Antibody Details

## Supplemental Materials

### Supplementary Figures

**Figure S1: Classification of ccRCC tissue immunophenotyped, example human tissue flow data cleanup, and marker MFI of human CD4+ Tconv cells.**

**Figure S2: Example mouse tissue flow data cleanup and BMDC cytotoxicity assay.**

**Figure S3: Cell sorting strategy, *Foxp3* transcript counts, and senescence-associated gene set enrichment analysis.**

**Figure S4: Background signal of tdTomato in fate-mapping mice and preservation of fluorescent protein signals from different fixation and permeabilization methods.**

**Figure S5: Function validation of** α**KLRG1-STR and** α**KLRG1-DLE, Treg expression of KLRG1, and therapeutic evaluation in MCA205 tumors**

## Acknowledgments

We want to thank Dr. Laurent Brossay, Dr. Eric Vivier and Innate Pharma for kindly sharing the anti-KLRG1 variable region sequences we adopted in this study. We also want thank Dr. Khashayarsha Khazaie and Dr. Fotini Gounari for providing mice and cell lines utilized in this study. Most importantly, we thank the patients and their families for their willing participation in this study.

## Funding

National Institutes of Health grant F32CA294616 (CJ)

Mayo Clinic Comprehensive Cancer Center: Gosselin Fellowship in Urologic Cancer and related Diseases of Males (CJ)

National Institutes of Health grant R01CA311117 (CRA)

National Institutes of Health grant R01CA300454-A1 (CRA)

Arizona Biomedical Research Centre (ABRC)

New Investigator Award grant RFGA2023-008-25 (CRA)

Arizona Biomedical Research Centre (ABRC)

New Investigator Award grant RFGA2026-03-011 (CRA)

Mayo Clinic Presidential Funds (CRA)

Crohn’s and Colitis Career Development Award #1045700 (FHH)

Mayo Clinic Comprehensive Cancer Center: Vranos Family Foundation Career Development Award in Prostate Cancer (JRA)

Mayo Clinic Comprehensive Cancer Center: Marley Career Development Award in Prostate Cancer (JRA)

Mayo Clinic Department of Immunology (CRA)

Mayo Clinic Department of Urology (CRA)

Mayo Clinic Genitourinary Disease Group (CRA)

## Author contributions

Conceptualization: CJ, CRA

Methodology: CJ, CRA

Formal analysis: CJ, CRA

Investigation: CJ, KRC, JCH, JFC, NB, PN, FS, NA

Resources: CJ, MMC, JNL, FHH, Y Z, PEA, HMA, CRA

Data curation: CJ, CRA

Writing - original draft: CJ, CRA

Writing - review & editing: CJ, CRA

Visualization: CJ, CRA

Supervision: CJ, CRA

Project administration: CJ, JRA, CRA

Funding acquisition: CJ, FHH, JRA, CRA

## Data and materials availability

RNAseq data will be deposited to GEO by time of publication. Flow cytometry data will be uploaded to Zenodo by time of publication. No software or algorithm was generated in this study. Any additional information required to reanalyze the data reported in this paper is available from the corresponding author upon reasonable request.

