## Supplemental Figures for "Cytotoxic CD4+ exTregs expressing KLRG1 retain immunosuppressive function in cancer"

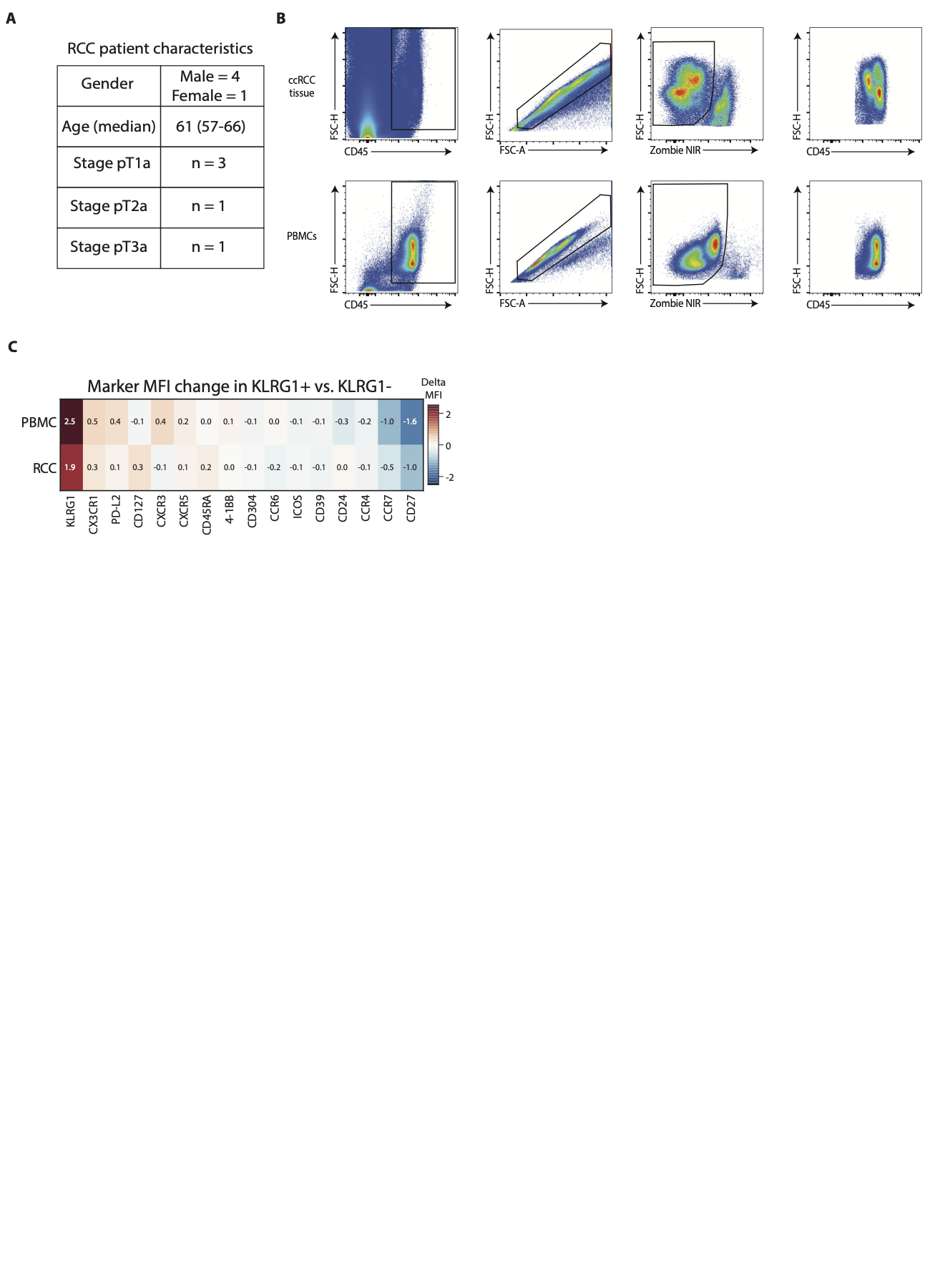

**Supplemental figure 1: Classification of ccRCC tissue immunophenotyped, example human tissue flow data cleanup, and marker MFI of human CD4+ Tconv cells**

(**A**) Patient characteristics of ccRCC tissue used in this study. (**B**) Representative pseudocolor cleaning gates for human ccRCC tissue (top) and cryopreserved human PBMCs (bottom) used throughout the study. (**C**) The delta MFIs for each significant marker from Figure 6I ccRCC or PBMCs. Cell color indicates whether the delta MFI is positive (red; up in KLRG1+ Tconv) or negative (blue; down in KLRG1+ Tconv).

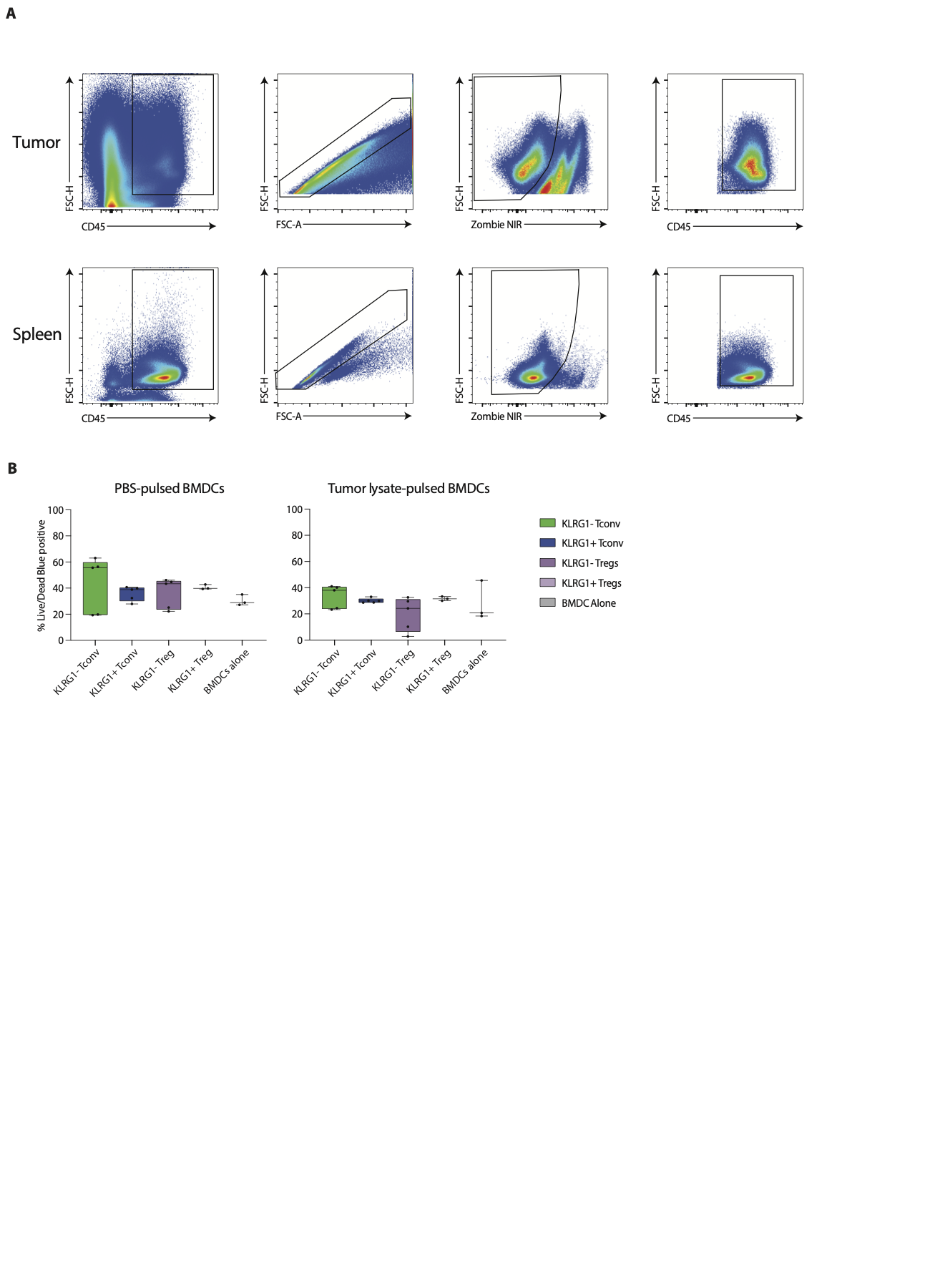

**Supplemental figure 2: Example mouse tissue flow data cleanup and BMDC cytotoxicity assay**

(**A**) Representative pseudocolor cleaning gates for mouse tumor tissue (top) and mouse splenic tissue (bottom) used throughout the study. (**B**) Frequency of Live/Dead blue positive BMDCs as a fraction of the total BMDC population after being pulsed overnight with PBS or MCA205 tumor lysate. Data shown are pooled data from two independent experiments with 1-3 technical replicates each.

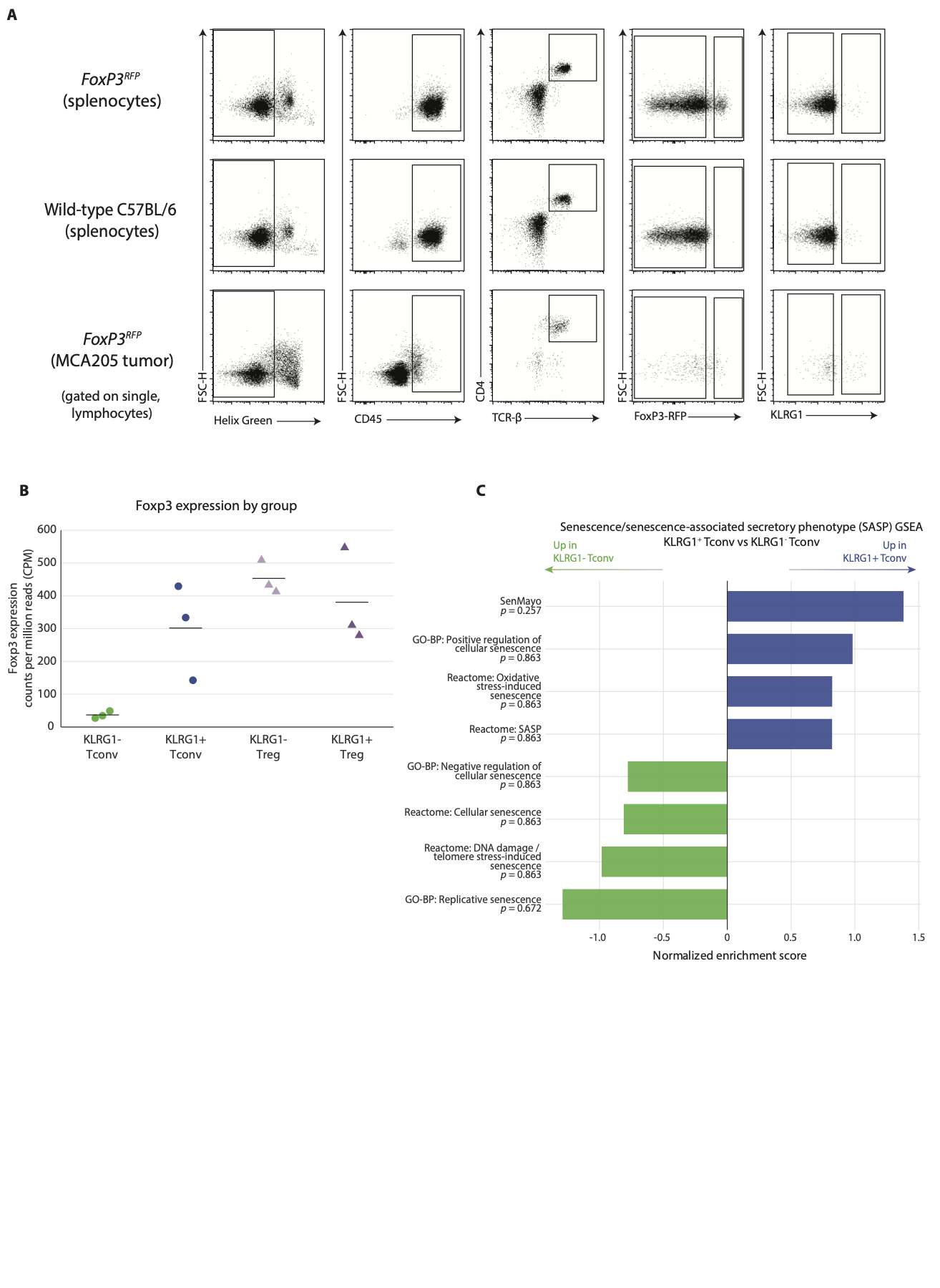

**Supplemental figure 3: Cell sorting strategy, *Foxp3* transcript counts, and senescence-associated gene set enrichment analysis**

(**A**) Representative plots of the gating strategy used to sort KLRG1+ and KLRG1- Tconv and Tregs from *Foxp3^RFP^* reporter mice for *in vitro* experiments in Figure 2 and RNA sequencing in Figure 4. Splenocytes from *Foxp3^RFP^* mice (top row) and from wild-type mice (middle row) were used to validate the RFP reporter and to set conservative gates on tumors (bottom row). (**B**) Raw *Foxp3* gene counts normalized to counts per one million reads in each CD4+ population. (**C**) Senescence and senescence-associated secretory phenotype (SASP) gene set enrichment analysis normalized enrichment scores. Gene set name and the associated BH-corrected *p*-value are listed along the left y-axis.

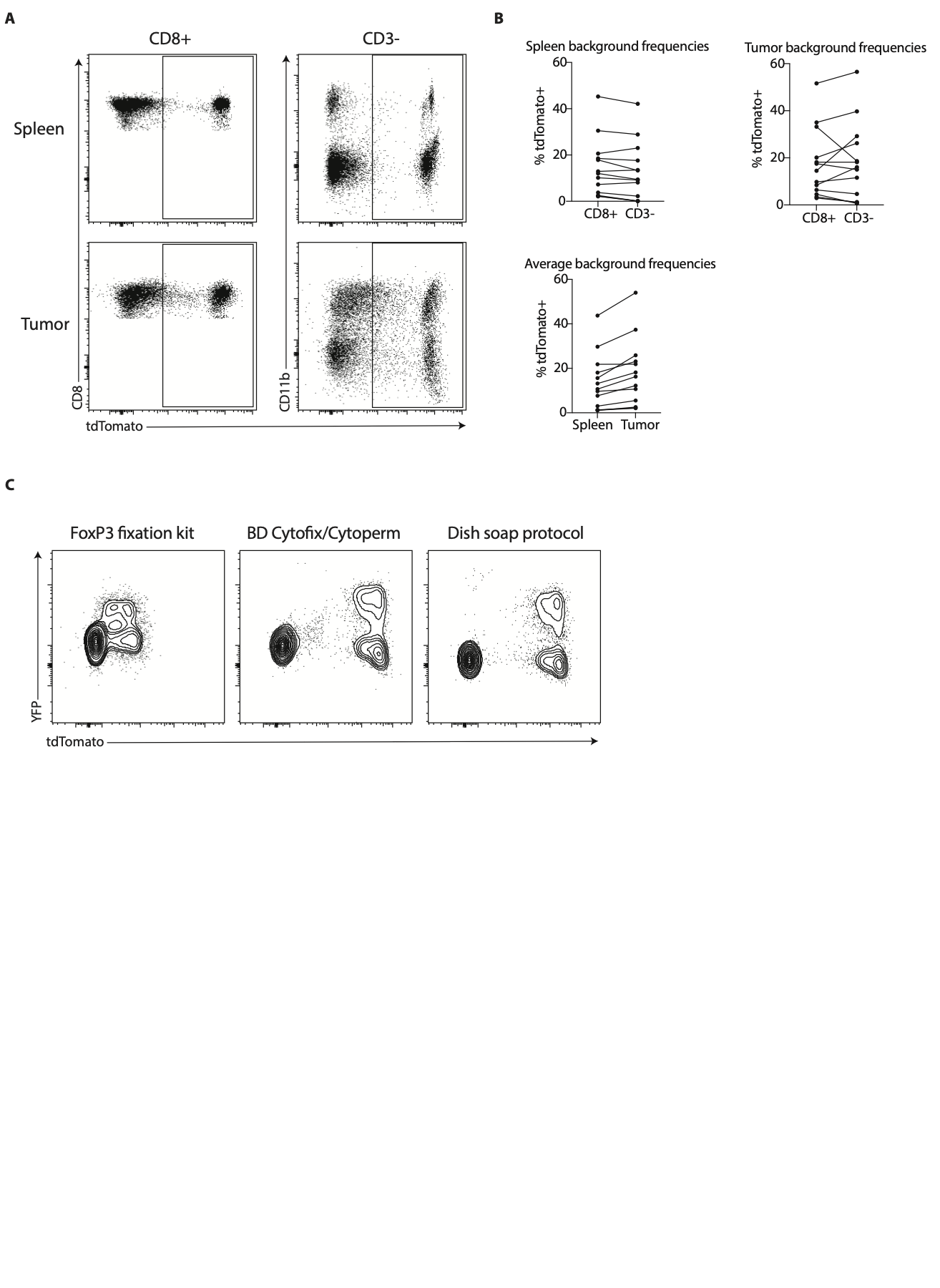

**Supplemental figure 4: Background signal of tdTomato in fate-mapping mice and preservation of fluorescent protein signals from different fixation and permeabilization methods**

(**A**) Representative plots of tdTomato signal in CD8+ T cells (left plots) and CD3- cells (right plots) that includes CD11b+ and CD11b- cells. Cells were gated on live CD45+ cells. Splenocytes represent the top row and tumor-derived cells represent the bottom row. (**B**) Quantitation of tdTomato+ cell frequency in CD8+ and CD3- cells from spleen (top left) and tumor tissue (bottom left). The average of CD8+ and CD3- tdTomato frequency in both spleen and tumor tissue (middle right). Lines are drawn between paired samples. Data shown include pooled data from groups of *n* = 6-7 mice from two independent experiments. (**C**) Representative comparison plots from splenocytes from *Foxp3*^YFP-Cre^ x *Rosa26*^CAG-LSL-TdTomato^ fate mapping mice using the eBioscience™ FoxP3/Transcription Factor Fixation/ Permeabilization kit (left), the BD Cytofix/Cytoperm™ Fixation/Permeabilization kit (center), and the dish soap protocol fixation/permeabilization method^30^. Plots give a qualitative view of fluorescent reporter signals YFP and tdTomato across each method.

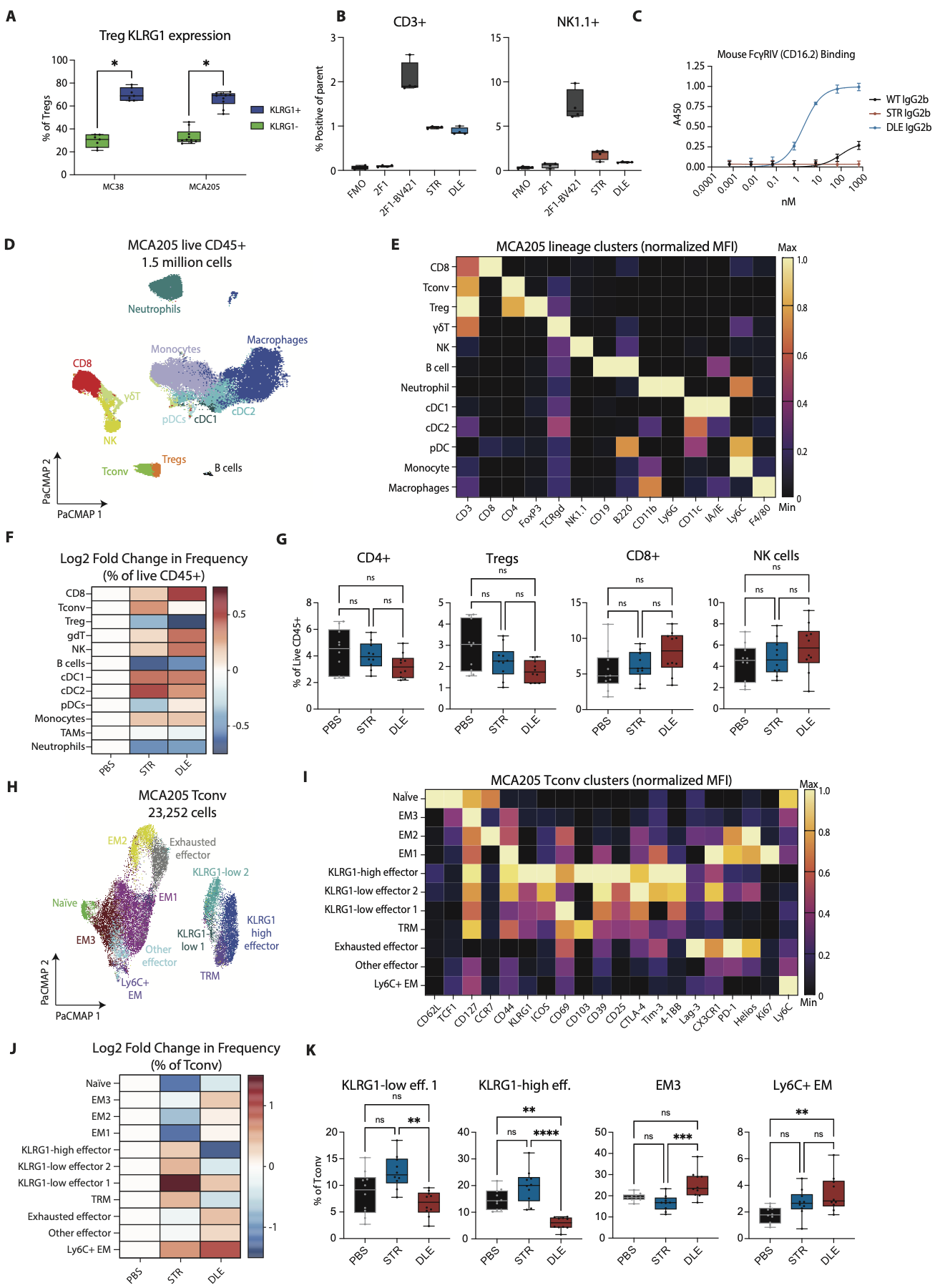

**Supplemental figure 5: Function validation of αKLRG1-STR and αKLRG1-DLE, Treg expression of KLRG1, and therapeutic evaluation in MCA205 tumors**

(**A**) Quantitation of KLRG1+ cells in a competitive binding flow cytometry assay. KLRG1+ cell frequency was assessed in CD3+ and NK1.1+ cells from mouse splenocytes. Data shown are pooled data from two independent experiments with *n* = 2 technical replicates each. FMO = sample stained with KLRG1-BV421; 2F1 = sample pre-incubated with unconjugated anti-mouse KLRG1 clone 2F1; 2F1-BV421 = sample with PBS vehicle as the pre-incubation reagent; STR = sample preincubated with in-house produced anti-KLRG1-STR; DLE= sample preincubated with in-house produced anti-KLRG1-DLE. (**B**) Sandwich ELISA quantitation of binding of in-house produced anti-KLRG1-STR, in-house produced anti-KLRG1-DLE, and a control WT IgG2b to mouse CD16.2. Data shown are pooled data from two independent experiments with 3 technical replicates each. Means are plotted with SEM shown. (**C**) Quantitation of tumor-infiltrating CD4+ Treg KLRG1 expression in MC38 and MCA205 tumors as fractions of Treg cells defined as FoxP3+ cells. Wilcoxon matched-pairs signed rank test was performed with Holm-Sidak multiple testing correction. (**D**) PaCMAP of live CD45+ cells from PBS, aKLRG1-STR, and aKLRG1-DLE-treated MC205 tumors. Clusters are derived from FlowSOM. (**E**) Normalized heatmap of lineage-defining marker MFI across all live CD45+ cells. (**F**) Log2 fold change of each lineage cluster frequency in each treatment arm, normalized to the frequency in the PBS group. (**G**) Frequency quantitation CD4+ T cells (left), Tregs (left-center), CD8+ T cells (right-center), and NK cells (right) as fractions of the live CD45+ clusters from each treatment group. Kruskal-Wallis one-way ANOVA with Dunn’s multiple testing correction was used for statistical testing. Data shown are from one experiment. All groups are *n* = 10. (**H**) PaCMAP plot of the CD4+ Tconv cluster from D. Clusters are derived from FlowSOM. (**I**) Normalized heatmap of phenotyping marker MFI in CD4+ Tconv across all treatment groups. (**J**) Log2 fold change of each CD4+ Tconv cluster frequency in each treatment arm, normalized to the frequency in the PBS group. (**K**) Frequency quantitation KLRG1-low effectors (eff.; left), KLRG1-high eff. (left-center), effector memory cluster 3 (EM3; right-center), and Ly6C+ effector memory (right) as fractions of the CD4+ Tconv clusters from each treatment group. Kruskal-Wallis one-way ANOVA with Dunn’s multiple testing correction was used for statistical testing. Data shown are from one experiment. All groups are *n* = 10. * = *p* < 0.05
